# New genetic signals for lung function highlight pathways and pleiotropy, and chronic obstructive pulmonary disease associations across multiple ancestries

**DOI:** 10.1101/343293

**Authors:** Nick Shrine, Anna L Guyatt, A Mesut Erzurumluoglu, Victoria E Jackson, Brian D Hobbs, Carl Melbourne, Chiara Batini, Katherine A Fawcett, Kijoung Song, Phuwanat Sakornsakolpat, Xingnan Li, Ruth Boxall, Nicola F Reeve, Ma’en Obeidat, Jing Hua Zhao, Matthias Wielscher, Understanding Society Scientific Group, Stefan Weiss, Katherine A Kentistou, James P Cook, Benjamin B Sun, Jian Zhou, Jennie Hui, Stefan Karrasch, Medea Imboden, Sarah E Harris, Jonathan Marten, Stefan Enroth, Shona M Kerr, Ida Surakka, Veronique Vitart, Terho Lehtimäki, Richard J Allen, Per S Bakke, Terri H Beaty, Eugene R Bleecker, Yohan Bossé, Corry-Anke Brandsma, Zhengming Chen, James D Crapo, John Danesh, Dawn L DeMeo, Frank Dudbridge, Ralf Ewert, Christian Gieger, Amund Gulsvik, Anna L Hansell, Ke Hao, Josh D Hoffman, John Hokanson, Georg Homuth, Peter K Joshi, Philippe Joubert, Claudia Langenberg, Xuan Li, Liming Li, Kuang Lin, Lars Lind, Nick Locantore, Jian’an Luan, Anubha Mahajan, Joseph C Maranville, Alison Murray, David C Nickle, Richard Packer, Margaret M Parker, Megan L Paynton, David Porteous, Dmitry Prokopenko, Dandi Qiao, Rajesh Rawal, Heiko Runz, Ian Sayers, Don D Sin, Blair H Smith, María Soler Artigas, David Sparrow, Ruth Tal-Singer, Paul RHJ Timmers, Maarten Van den Berge, John C Whittaker, Prescott Woodruff, Laura M Yerges Armstrong, Olga G Troyanskaya, Olli T Raitakari, Mika Kähönen, Ozren Polasek, Ulf Gyllensten, Igor Rudan, Ian J Deary, Nicole M Probst-Hensch, Holger Schulz, Alan L James, James F Wilson, Beate Stubbe, Eleftheria Zeggini, Marjo-Riitta Jarvelin, Nick Wareham, Edwin K Silverman, Caroline Hayward, Andrew P Morris, Adam S Butterworth, Robert A Scott, Robin G Walters, Deborah A Meyers, Michael H Cho, David P Strachan, Ian P Hall, Martin D Tobin, Louise V Wain

**Author notes:** = Contributed equally to this work (Nick Shrine, Anna L Guyatt, A Mesut Erzurumluoglu, Ian P Hall, Martin D Tobin, Louise V Wain). = Corresponding Authors.

## Abstract

Reduced lung function predicts mortality and is key to the diagnosis of COPD. In a genome-wide association study in 400,102 individuals of European ancestry, we define 279 lung function signals, one-half of which are new. In combination these variants strongly predict COPD in deeply-phenotyped patient populations. Furthermore, the combined effect of these variants showed generalisability across smokers and never-smokers, and across ancestral groups. We highlight biological pathways, known and potential drug targets for COPD and, in phenome-wide association studies, autoimmune-related and other pleiotropic effects of lung function associated variants. This new genetic evidence has potential to improve future preventive and therapeutic strategies for COPD.

## Introduction

Impaired lung function is predictive of mortality^1^ and is the key diagnostic criterion for chronic obstructive pulmonary disease (COPD). Globally, COPD accounted for 2.9 million deaths in 2016^2^, being one of the key causes of both Years of Life Lost and Years Lived with Disability worldwide^3^. Determinants of maximally attained lung function and of lung function decline can influence the risk of developing COPD. Tobacco smoking is the single largest risk factor for COPD, although other environmental exposures and genetic makeup are important^4,5^. Genetic variants associated with lung function and COPD susceptibility can be causally informative, assisting with risk prediction, as well as drug target identification and validation^6^. Whilst there has been considerable progress in identifying genetic markers associated with lung function and risk of COPD^4,7-19^ seeking a high yield of associated genetic variants is key to progressing knowledge because: (i) implication of multiple molecules in each pathway will be needed to build an accurate picture of the pathways underpinning development of COPD; (ii) not all proteins identified will be druggable and; (iii) combining information across multiple variants can improve prediction of disease susceptibility.

Through new detailed quality control and analyses of spirometric measures of lung function in UK Biobank, completion of genome-wide genotyping in UK Biobank, and expansion of the SpiroMeta Consortium, we undertook the largest genome-wide association study of lung function performed to date. Comprising a total of 400,102 individuals of European ancestry, our study entailed a near seven-fold increase in sample size over previous studies of similar ancestry to address the following aims: (i) to generate a high yield of genetic markers associated with lung function; (ii) to confirm and fine-map previously reported lung function signals; (iii) to investigate the putative causal genes and biological pathways through which lung function associated variants act, and their wider pleiotropic effects on other traits; and (iv) to generate a weighted genetic risk score for lung function and test its association with COPD susceptibility in individuals of European and other ancestries.

## Results

### 139 new signals for lung function

Here we present a total of 279 distinct association signals for lung function, of which a half (139 variants) are new having reached genome-wide significance (P<5×10^−9^) in this study. We increased the sample size available for the study of quantitative measures of lung function in UK Biobank by refining the quality control of spirometry based on recommendations of the UK Biobank Outcomes Adjudication Working Group, utilising additional metrics derived from the blow curve time series measurements, and relaxing the reproducibility threshold for repeat measures (**Supplementary Note**). Genome-wide association analyses of forced expired volume in 1 second (FEV_1_), forced vital capacity (FVC) and FEV_1_/FVC were then undertaken in 321,047 individuals in UK Biobank (**Supplementary Table 1**) and in 79,055 individuals from the SpiroMeta Consortium (**Supplementary Tables 2 and 3**). A linear mixed model approach implemented in BOLT-LMM^20^ was used for UK Biobank to account for relatedness and fine-scale population structure (**Online Methods**). A total of 19,871,028 variants imputed in both UK Biobank and SpiroMeta were analysed. Peak expiratory flow (PEF) was also analysed genome-wide in UK Biobank and up to 24,218 samples from SpiroMeta. All individuals included in the genome-wide analyses were of European ancestry (**Supplementary Figure 1** and **Supplementary Table 2**).

To maximise statistical power for discovery of new signals, whilst maintaining stringent significance thresholds to minimise reporting of false positives, we adopted a study design incorporating both two-stage and one-stage approaches (**Figure 1**). In the two-stage analysis, 99 new distinct signals, defined using conditional analyses, were associated with one or more traits at P<5×10^−9^ in UK Biobank and showed association (P<10^−3^) with a consistent direction of effect in SpiroMeta (“Tier 1” signals, **Supplementary Figure 2; Supplementary Table 4**). In the one-stage analysis, we meta-analysed UK Biobank and SpiroMeta (up to 400,102 individuals) and 40 additional new distinct signals associated with one or more lung function traits reaching P<5×10^−9^ were identified (**Supplementary Figure 2, Supplementary Table 4**) that were also associated with P<10^−3^ separately in UK Biobank and in SpiroMeta, with consistent direction of effect (“Tier 2” signals). An additional 323 signals were significantly associated with one or more lung function traits in the meta-analysis of UK Biobank and SpiroMeta (P<5×10^−9^) and reached P<10^3^ for association in only one of UK Biobank or SpiroMeta (“Tier 3” signals, **Supplementary Table 5**). Only the 139 signals meeting Tier 1 and Tier 2 criteria were followed up further. The strength and direction of association of the sentinel variant (the variant in each signal with the lowest P value) for these 139 new signals across all 4 lung function traits are shown in **Figure 2**.

**Figure 1:**
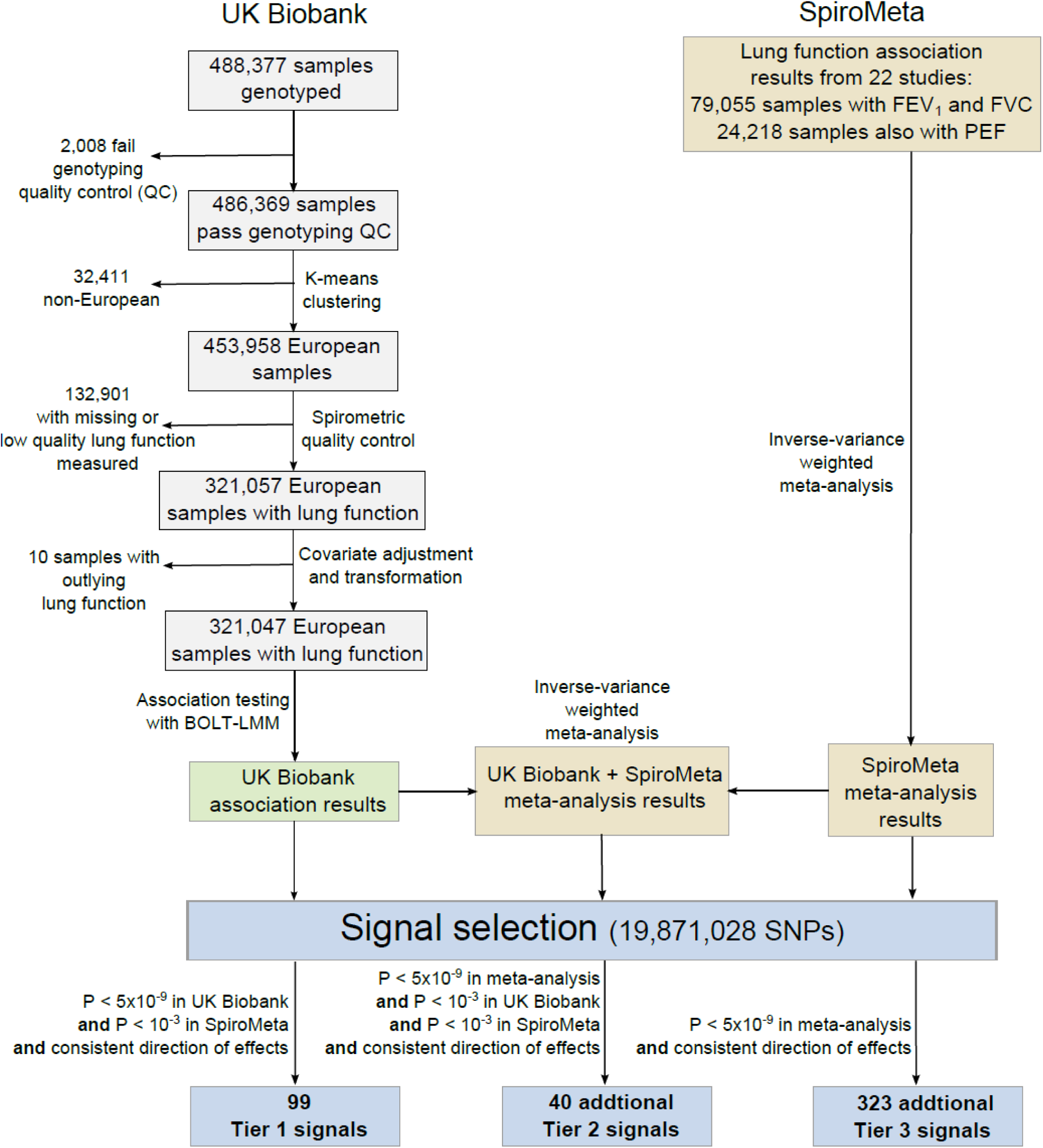
Study design. Tier 1 signals had P<5×10^−9^ in UK Biobank and P<10^−3^ in SpiroMeta with consistent direction of effect. Tier 2 signals had P<5×10^−9^ in the meta-analysis of UK Biobank and SpiroMeta with P<10^−3^ in UK Biobank and P<10^−3^ in SpiroMeta with consistent directions of effect. Signals with P<5×10^−9^ in the meta-analysis of UK Biobank and SpiroMeta, and that had consistent directions of effect but did not meet P<10^−3^ in both cohorts were reported as Tier 3.

**Figure 2:**
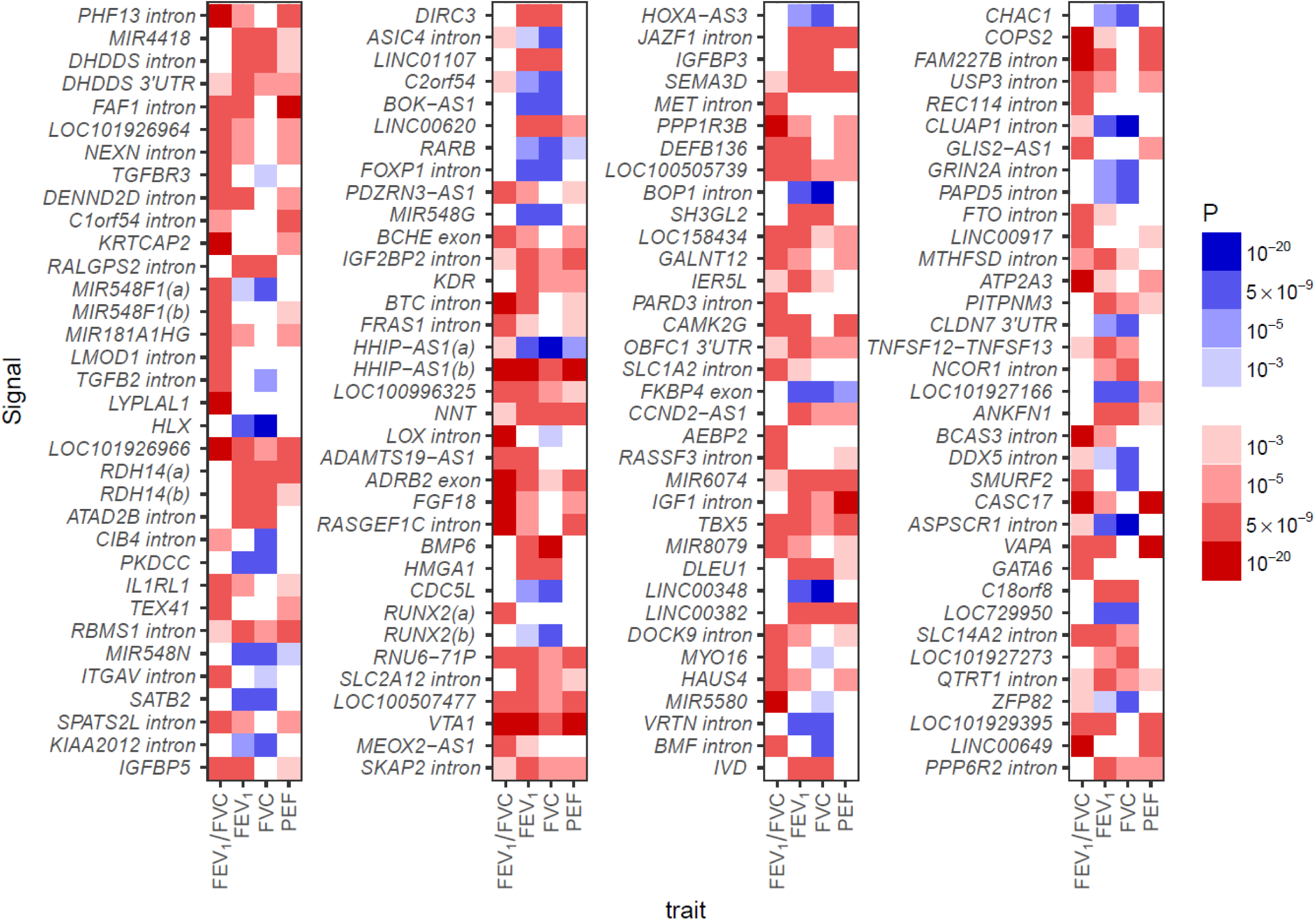
Strength and direction of association across four lung function traits for 139 novel signals: Red indicates decrease in the lung function trait; blue indicates an increase. All effects are aligned to the allele associated with decreased FEV_1_/FVC, hence the FEV_1_/FVC column is only red or white. P-values are from the meta-analysis of UK Biobank and SpiroMeta (n=400,102). The scale points are thresholds used for (i) confirmation in 2-stage analysis and 1-stage analysis (P<10^−3^); (ii) confirmation of association of previous signals (P<10^−5^); (iii) signal selection in 2-stage and 1-stage analysis (P<5×10^−9^); capped at (P<10^−20^).

To assess whether any of these 139 signals associated with lung function could be driven via an underlying association with smoking, we examined association of the sentinel variants with smoking behaviour in UK Biobank (**Online Methods**). The only new sentinel associated with smoking behaviour was rs193686 (in an intron of *MET*, **Supplementary Table 6**). Therefore, we tested for association between this variant and lung function in never smokers (n=173,658). Whilst rs193686 was associated with smoking initiation (P=9.18×10^−6^), the allele associated with smoking initiation was associated with increased lung function in never smokers (FEV_1_/FVC P=5.28×10^−10^, **Supplementary Table 7**). Therefore, this signal was retained for further analysis.

### A total of 279 signals of association for lung function

Of 157 previously published signals of association with lung function and COPD^3,6–18^, 142 were associated at P<10^−5^ in UK Biobank (**Online Methods, Supplementary Figure 3**, **Supplementary Table 8**). Two sentinel variants (rs1689510 near *RAB5B* and rs11134789 in an intron of *ADAM19*) were associated with smoking initiation (P=9.72×10^−6^ and P=2.13×10^−5^, respectively) (**Supplementary Table 6**), but were also associated with lung function in never smokers (P=2.49×10^−8^ for FEV_1_ and P=2.94×10^−45^ for FEV_1_/FVC, respectively, **Supplementary Table 7**). SNP rs17486278 at *CHRNA5* and rs11667314 near *CYP2A6* were each associated with cigarettes per day (P=1.35×10^−79^ and P=6.47×10^−24^, respectively; **Supplementary Table 6**); neither were significantly associated with lung function among never smokers, hence these latter two signals were excluded from further analysis. This brings the total number of distinct signals of association with lung function to 279 (**Supplementary Table 9**). None of these variants showed interaction with ever-smoking status (P>1.8×10^−4^, Online Methods**, Supplementary Table 7**). The 140 previously reported lung function signals showing association in this study (UK Biobank P<10^−5^) explained 5.0%, 3.4%, 9.2% and 4.5% of the estimated heritability of FEV_1_, FVC, FEV_1_/FVC and PEF, respectively (**Online Methods**). The 139 new signals reported here, explain an additional 4.3%, 3.3%, 3.9% and 3.3% of the estimated heritability, respectively.

### Identification of putative causal genes

Bayesian refinement was undertaken for each signal to identify the set of variants that were 99% likely to contain the underlying causal variant (assuming the causal variant has been analysed). The signals in the HLA region were excluded due to extended linkage disequilibrium. The results from the meta-analysis of UK Biobank and SpiroMeta were used to define the 99% credible sets (**Online Methods**, **Supplementary Table 10, Supplementary File–Region Plots**).

To identify putative causal genes for each signal, we identified deleterious variants and variants associated with gene expression (eQTLs) or protein levels (pQTLs) within each 99% credible set for all new and previously reported signals outside the HLA region (**Online Methods**).

There were 25 SNPs, located in 22 unique genes, which were exonic, at a splice site or in the untranslated regions and additionally annotated as potentially deleterious (**Online Methods**, **Supplementary Table 11**). Amongst our new signals, there were 10 variants annotated as deleterious in 9 different genes: *DOCK9* (rs117633128, MAF=10.6%), *CEP72* (rs12522955, MAF=20.2%), *BCHE* (rs1799807, MAF=1.95%), *DST* (rs11756977, MAF=28.9%), *KIAA0753* (rs2304977, MAF=37.7%; rs9889363, MAF=37.7%), *LRRC45* (rs72861736, MAF=10.9%), *BTC* (rs11938093, MAF=26.6%), *C2orf54* (rs6709469, MAF=49.9%) and *IER5L* (rs184457, MAF=31.5%). Of these, the missense variant in *BCHE* (rs1799807) had the highest posterior probability (0.996) in its respective credible set, was low frequency (MAF=1.95%) and resulted in an amino acid change from aspartic acid (D) to glycine (G), known to affect the function of the encoded butyrylcholinesterase enzyme by altering substrate binding^21^. The two common missense variants in *KIAA0753* were within the credible set of new signal rs4796334. *KIAA0753*, *CEP72* and *LRRC45* all encode proteins with a role in ciliogenesis or cilia maintenance^22–26^, and all are highly expressed in the airway epithelium^27^.

Variants in the 99% credible sets (n=9,698) were queried in three eQTL resources to identify associations with gene expression in lung^28–30^ (sample size n=1,111; **Supplementary Table 12**), blood^31^ (n=4,896) and a subset of GTEx^32^ tissues (max n=388, **Online Methods**). The tissues included from GTEx were lung and blood, plus nine tissues known to contain smooth muscle (**Online Methods**). The latter were chosen based on previous reports of enrichment of lung function GWAS signals in smooth muscle-containing tissues^18,33^. We identified 88 genes for which the most significant SNP associated with expression of that gene in the respective eQTL resource was within one of the 99% credible sets. These 88 genes were implicated by 58 of the 279 signals (**Supplementary Table 13**).

We checked credible set variants for association with protein levels in a pQTL study^34^ comprising SNP associations for 3,600 plasma proteins. Using a Bonferroni-corrected 5% significance threshold for 276 tests for these 3,600 proteins (P < 5.03×10^−8^), we found 1,076 pQTLs in our credible sets covering 26 lung function sentinels implicating 34 proteins. For 5 of these proteins the pQTL sentinel was contained within our lung function credible set: ECM1, THBS4, NPNT, C1QTNF5 and SCARF2 (**Supplementary Table 14**).

In total, 107 putative causal genes were identified (**Table 1**), 8 by both a deleterious variant and an eQTL signal (including *KIAA0753* implicated by two deleterious variants), 1 (*NPNT)* by both an eQTL and a pQTL signal, 1 (*SCARF2*) by both a deleterious variant and a pQTL signal, 13 by a deleterious variant only, 81 by an eQTL signal only and 3 by a pQTL signal only. Among these 107 genes, we highlight 75 for the first time as putative causal genes for lung function (43 implicated by a new signal and 32 newly implicated by a previous signal^18^).

**Table 1:**
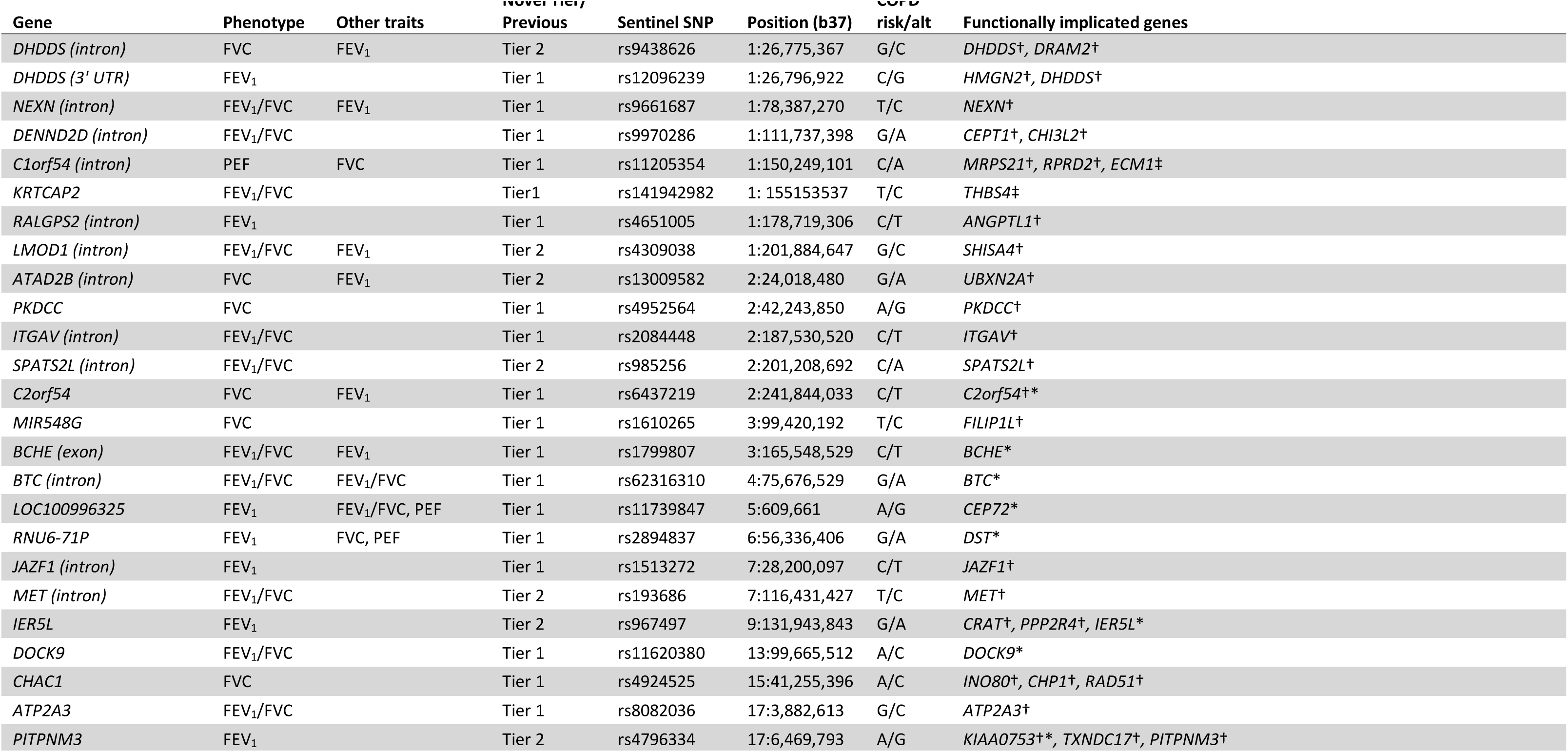
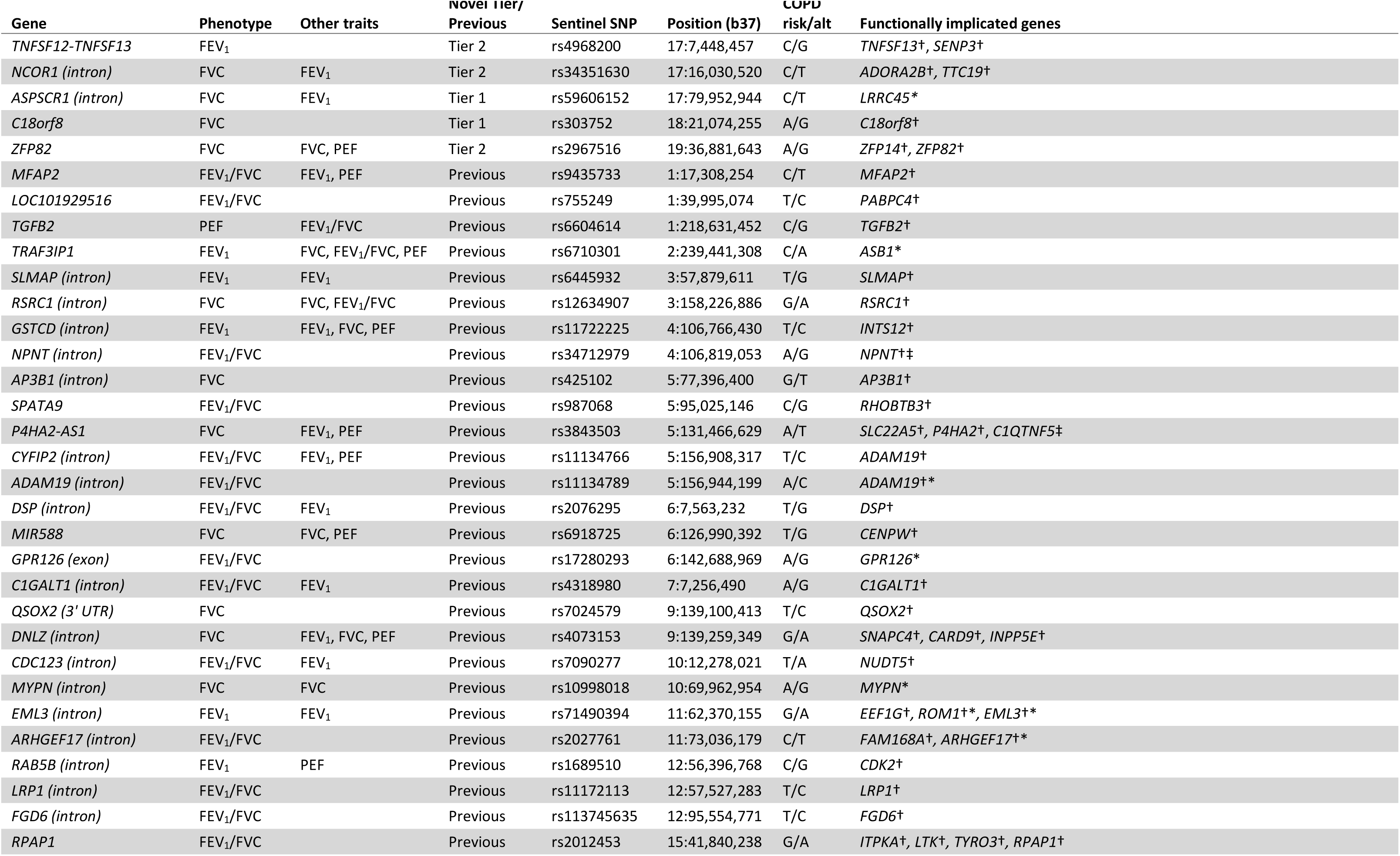
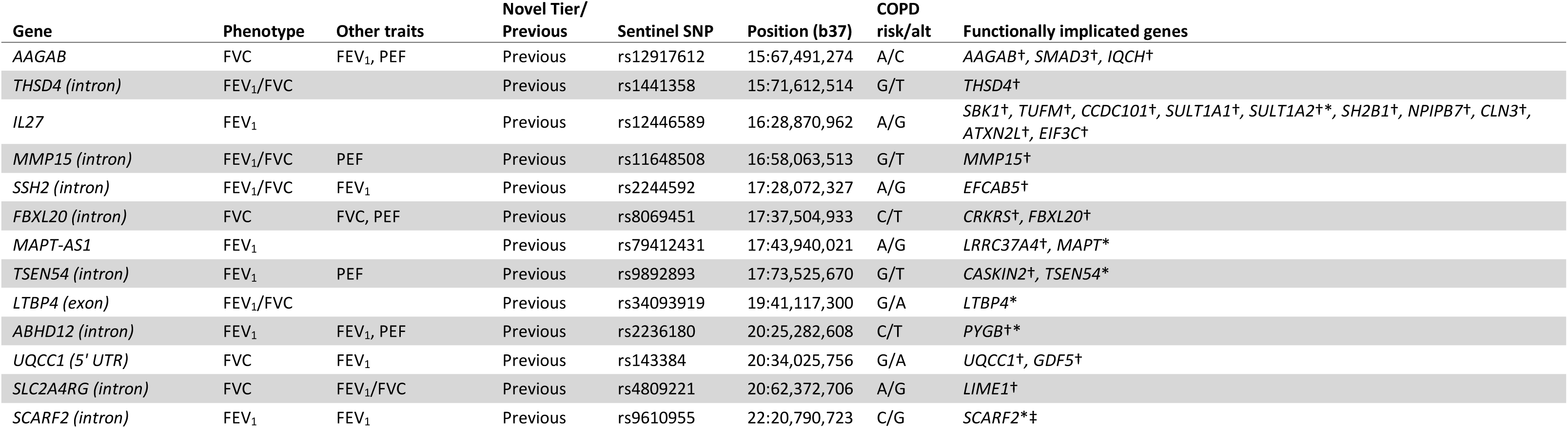
Genes implicated using gene expression data, protein level data and functional annotation. ^†^Genes implicated by eQTL signals: Lung eQTL (n=1,111) and Blood eQTL (n=4,896) datasets and eleven GTEx (V7) tissues were screened: Artery Aorta (n=267), Artery Coronary (n=152), Artery Tibial (n=388), Colon Sigmoid (n=203), Colon Transverse (n=246), Esophagus Gastroesophageal Junction (n=213), Esophagus Muscularis (n=335), Lung (n=383), Small Intestine Terminal Ileum (n=122), Stomach (n=237), and Whole Blood (n=369); see **Supplementary Table 13** for direction of gene expression for the COPD (lung function reducing) risk allele. ^‡^Genes implicated by pQTL signals: pQLT look up in 3,600 plasma proteins (n up to 3,300). *Genes implicated because they contain a deleterious variant (**Supplementary Table 11**). “Other traits” column lists the other lung function traits for which the sentinel was associated at P<5×10^−9^ in the meta-analysis of UK Biobank and SpiroMeta.

### Pathway analysis

We tested whether these 107 putative causal genes were enriched in gene sets and biological pathways (**Online Methods**), finding an enrichment of genes in elastic fibre and extracellular matrix organisation pathways, and a number of gene ontologies including gene sets relating to the cytoskeleton and processes involved in ciliogenesis (for example, cytoskeleton organisation, organelle organisation, centriole replication and microtubule-based processes) (**Supplementary Table 15**). Whilst the enrichment in elastic fibre-related pathways is consistent with our previous study^18^, enrichment in these pathways was further supported in this analysis by two new genes, *ITGAV* (at a new signal) and *GDF5* (a newly implicated gene for a previously reported signal), and by strengthened eQTL evidence for *TGFB2* and *MFAP2* as the putative causal genes at two previously reported signals. The presence of *TGFB2*, *GDF5* and *SMAD3* in our list of 107 genes resulted in enrichment of a TGF-β superfamily signalling pathway (TGF-Core) and multiple related gene ontology terms (**Supplementary Table 15**).

### Functional enrichment analyses

We tested for enrichment of the 279 lung function signals in DNase I hypersensitivity sites in 125 cell lines from ENCODE and 299 cell lines and tissues from RoadMap Epigenome Project using FORGE v1.1^35^. There was significant tissue specific overlap (**Online Methods**) of the 279 signals with DNAse1 hotspots in adult and foetal lung, foetal muscle (skeletal), foetal stomach, foetal heart, and fibroblasts (**Supplementary Figure 4**).

We used DeepSEA^36^, a variant effect predictor which utilises a deep-learning algorithm, to identify whether our signals were predicted to have a chromatin effect in lung-related cell lines. We identified 10 signals (including 5 new signals) for which the SNP with the largest posterior probability of being causal also had a significant predicted effect on a DNase I hypersensitivity site in lung-related cells (**Supplementary Table 16**). This included a new signal near *SMURF2* (17q24.1, rs11653958) that also had a predicted functional effect on histone marks (DNase I hypersensitivity sites, H3K9ac, H3K27ac, H3K4me1, H3K4me2, H3K4me3) and on *CEBPB*, *FOSL2*, *SIN2AK-20* and *TCF12* transcription factor binding sites, and a new signal near *PDZRN3-AS1* (rs586936) had a large predicted effect on a CEBPB transcription factor binding site.

### Drug targets

All 107 putative causal genes were interrogated against the gene-drug interactions table of the Drug-Gene Interactions Database (DGIDB)^37^ (**Supplementary Table 17**). We highlight two examples of new genetic signals implicating targets for drugs in development for indications other than COPD. One of our new signals is an eQTL for *ITGAV*. *ITGAV* encodes a component of the avβ6 integrin heterodimer, which is inhibited by a monoclonal antibody, STX-100, in development for pulmonary fibrosis (ClinicalTrials.gov Identifier: NCT01371305) and for which the small molecule GSK3008348 (ClinicalTrials.gov Identifier: NCT03069989) is an antagonist^38^. Integrins have an emerging role as local activators of TGFβ and specifically the avb6 integrin heterodimer can activate latent-TGFβ^39^. In our study, the allele associated with reduced expression of *ITGAV* (**Supplementary Table 13**) was associated with reduced risk of COPD (**Supplementary Table 9**) suggesting that inhibitors of αvβ6 integrin might also have a beneficial effect in COPD. Another of our new signals is associated with expression of *TNFSF13* (synonym *APRIL*), a cytokine which is a member of the TNF ligand family. Atacicept blocks B cell stimulation by *TNFSF13* (as well as by BLyS) and reduced systemic lupus erythematosus disease activity in a recent Phase IIb trial^40^. In our study, the allele associated with decreased expression of *TNFSF13* was associated with reduced FEV_1_, indicating that vigilance for pulmonary consequences of atacicept may be warranted.

### Genetic Risk Score: association with FEV_1_/FVC and COPD in multiple ancestries

We constructed a genetic risk score (GRS) weighted by FEV_1_/FVC effect sizes comprising all 279 new or previously reported sentinel variants, and tested the association of the GRS with FEV_1_/FVC and GOLD Stage 2-4 COPD (FEV_1_/FVC<0.7 and FEV_1_<80% predicted) in different ancestry groups in UK Biobank, and China Kadoorie Biobank (**Online Methods**, **Supplementary Table 18**). The GRS was associated with FEV_1_/FVC and COPD in each of the ancestry groups (**Figure 3A**).

**Figure 3:**
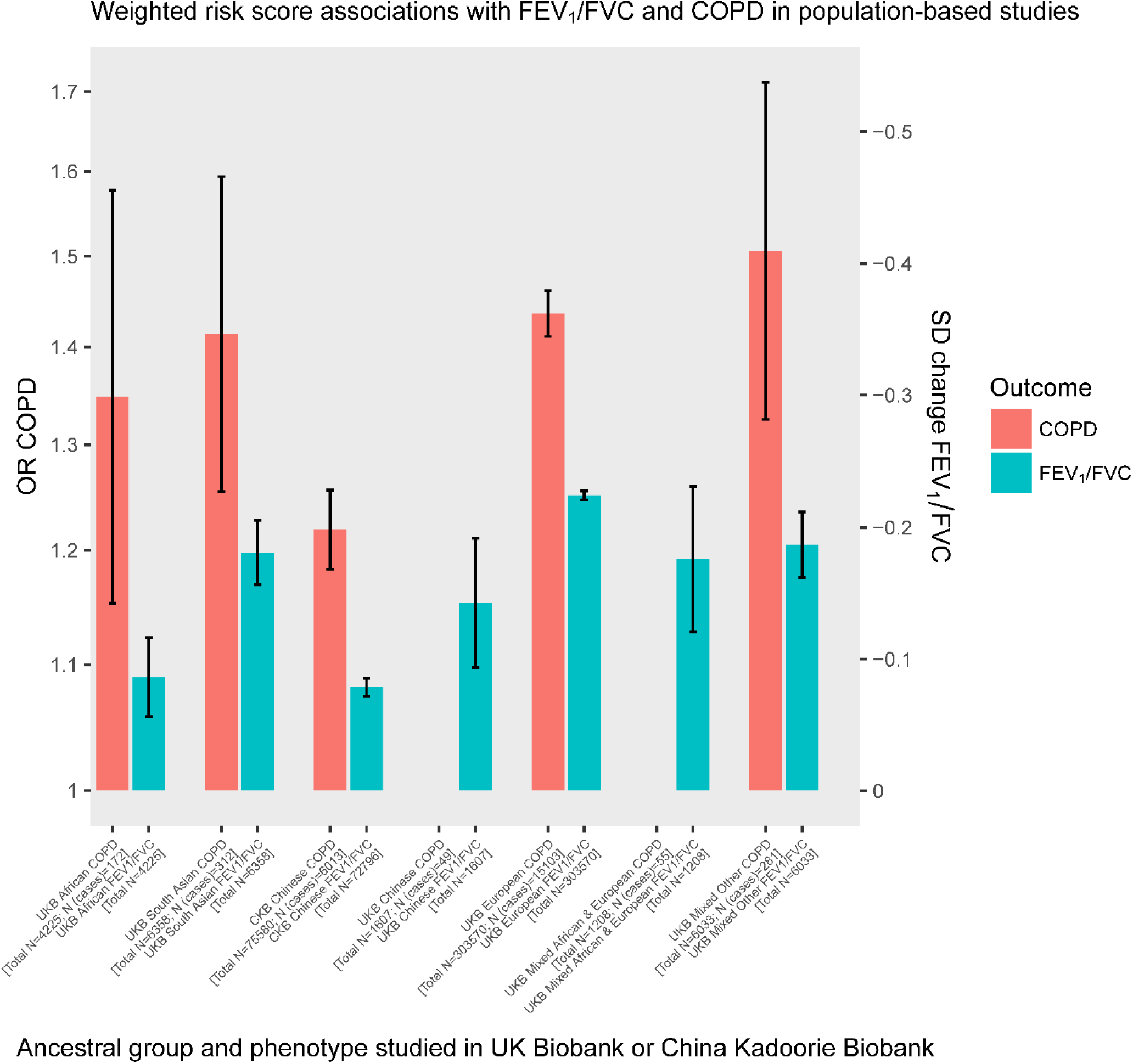
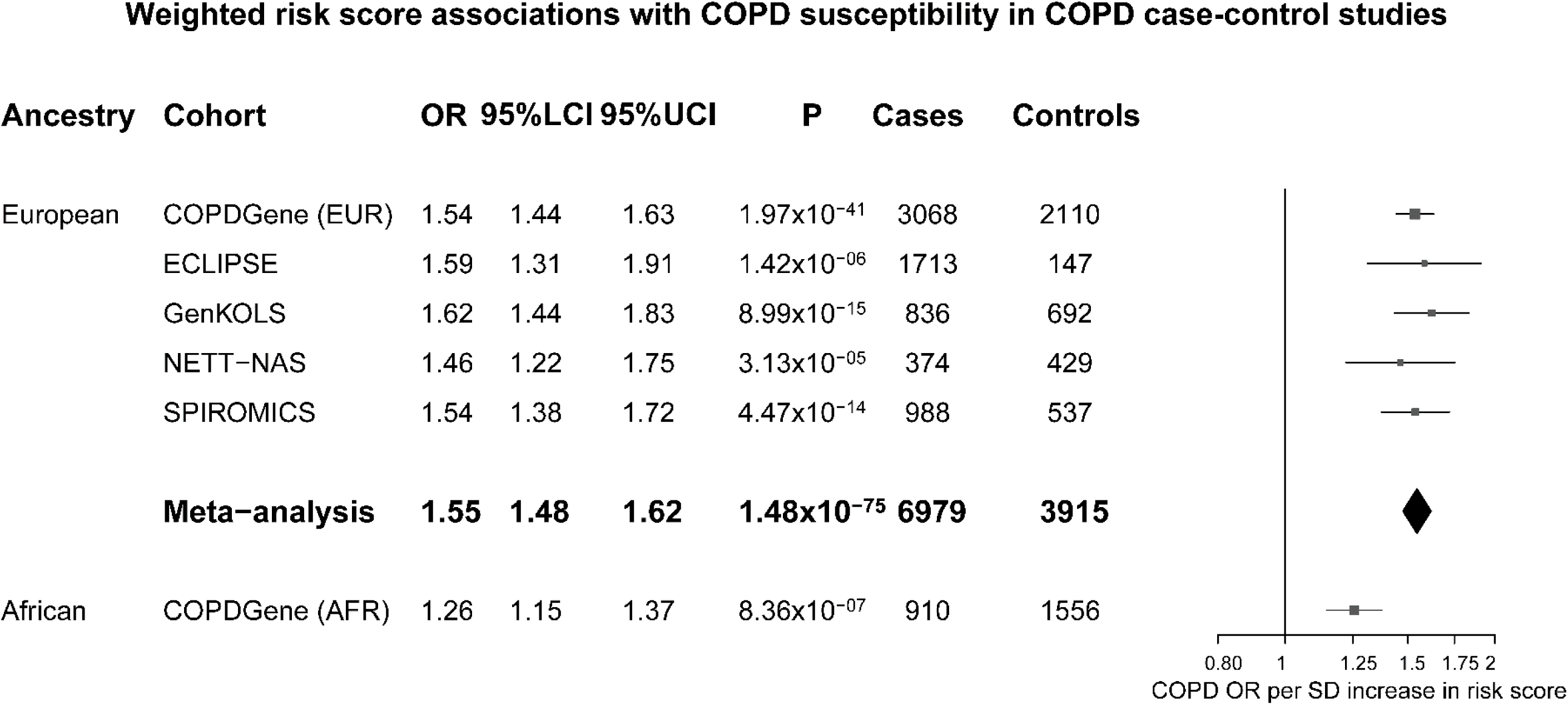
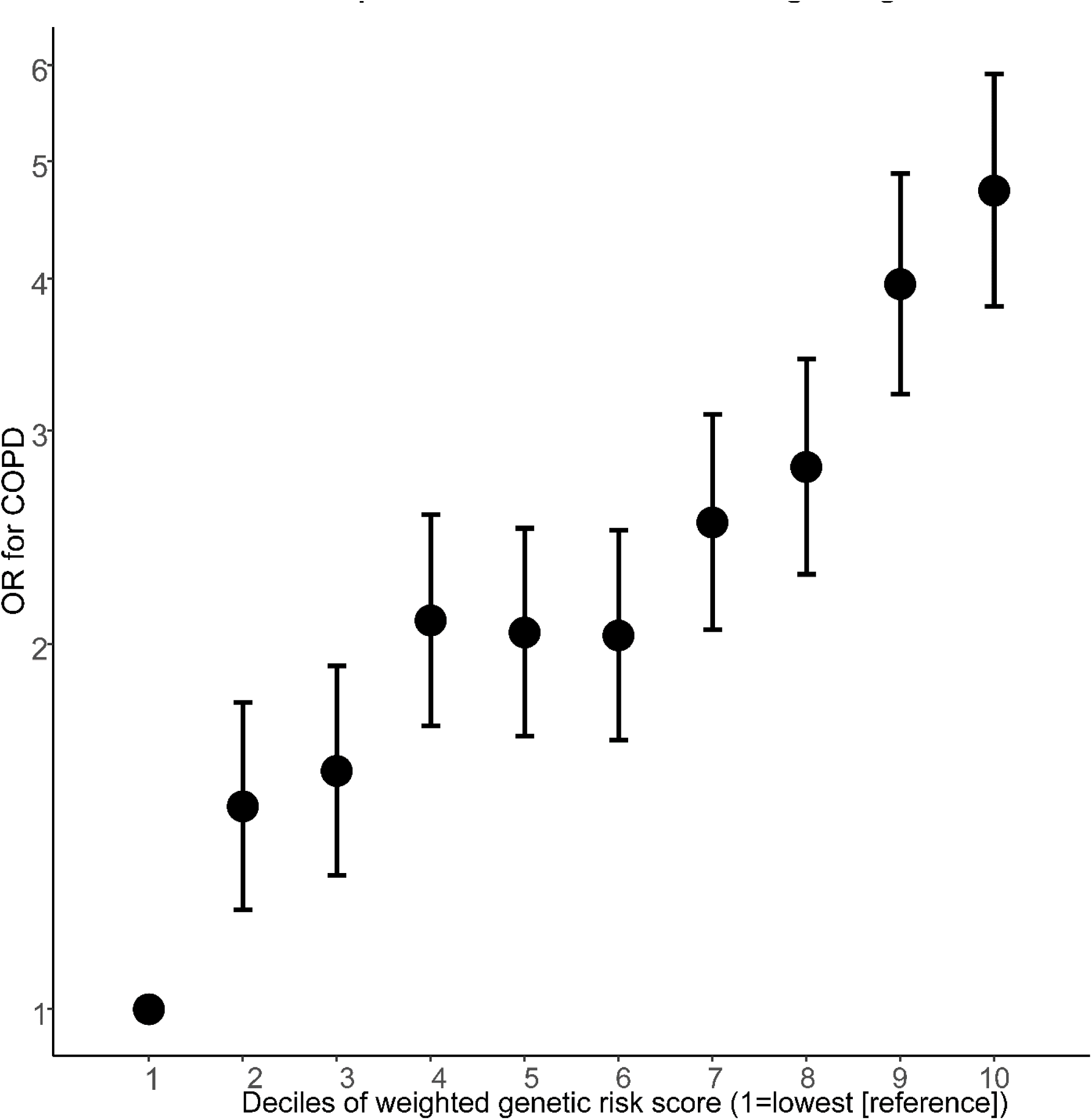
Association of weighted genetic risk score (GRS) with COPD and FEV_1_/FVC. **A.** Association of weighted genetic risk score (GRS) with COPD and FEV_1_/FVC in UK Biobank and China Kadoorie Biobank (CKB). The left axis denotes odds ratios (OR) for COPD per 1 standard deviation (SD) increase in weighted GRS (OR for COPD shown only for ancestries in UK Biobank with > 100 cases of COPD). COPD was defined as FEV_1_/FVC < 0.7 and FEV_1_ < 0.8 of the predicted value, i.e. GOLD stage 2-4 categorisation. Bars (in red) are labelled with ancestral groups, and the total sample size and number of COPD cases are given. The right-hand axis denotes change in standard deviation (SD) units of FEV_1_/FVC per 1 SD increase in weighted GRS in the same individuals (blue bars). For means and standard deviations of the risk scores in each group, see **Supplementary Table 18.** Note some variants featuring in the GRS were discovered in UK Biobank individuals of European ancestry. The height of the bars represents the effect estimate, and the black whiskers represent 95% confidence intervals. There were 13 SNPs with MAF <0.1% in at least one ancestral group: 13/279 in Chinese (of which 4/13 were monomorphic). Two of the 13 SNPs that were monomorphic in Chinese people had MAF<0.1% in Africans. **B.** Odds ratio (OR) for COPD per 1 standard deviation (SD) increase in weighted genetic risk score in each of six study groups (COPDGene [Non-Hispanic White], COPDGene [African-American], ECLIPSE, GenKOLS, SPIROMICS, NETT-NAS). COPD was defined using GOLD 2-4 criteria. For means and standard deviations of the risk scores in each group see **Supplementary Table 20**. The vertical black line indicates the null effect (an OR of 1). The point estimate of each study is represented by a box proportional to the study’s weight, with the lines representing the lower and upper bounds of the 95% confidence interval. A fixed effect meta-analysis of the five European-ancestry groups is denoted with a diamond, the width of which represents the 95% confidence interval for the estimate (I^2^ statistic=0). **C.** Odds ratios (OR) for COPD according to membership of deciles 2-10 of the weighted genetic risk score, with decile 1 as the reference group (the 10% of individuals with the lowest genetic risk score). Each point represents a meta-analysis of results for a given comparison (i.e. decile 2 vs reference, decile 3 vs reference … decile 10 versus reference) in five external European-ancestry study groups (COPDGene, ECLIPSE, GenKOLS, SPIROMICS, NETT-NAS). Deciles were calculated and models were run in each group separately. Points represent odds ratios, and error bars correspond to 95% confidence intervals (**Supplementary Table 21**).

We tested for a GRS interaction with smoking in European ancestry individuals in UK Biobank^41^. No statistical interaction was seen for FEV_1_/FVC (interaction term −0.002 per SD change in GRS, 95% CI: [0.009, 0.005], P=0.532), whilst the findings for COPD were consistent with a slightly smaller effect of the GRS in ever-smokers (OR for ever-smoking-GRS interaction term per SD change in GRS 0.96, 95% CI: [0.92, 0.99], P=0.015).

The association of the GRS with COPD susceptibility was additionally tested in deeply-phenotyped case-control studies (**Supplementary Table 19**). Similar effect size estimates were seen across each of the 5 European ancestry studies (**Figure 3B**); in the meta-analysis of these studies (n=6,979 cases and 3,915 controls), the odds ratio for COPD per standard deviation of the weighted GRS was 1.55 (95% CI: [1.48, 1.62]), P=2.87×10^−75^ (**Supplementary Table 20**). The GRS was also associated with COPD in individuals of African-American ancestry in COPDGene (P=8.36×10^−7^), albeit with a smaller effect size estimate, odds ratio=1.26 (95% CI: [1.15, 1.37]).

To aid clinical interpretation, we divided individuals in each of the European ancestry deeply-phenotyped COPD case-control studies into deciles, according to their value of the weighted GRS. The odds ratio for COPD in members of the highest GRS decile compared to the lowest GRS decile was 4.73 (95% CI: [3.79, 5.90]), P=3.00×10^−43^ (**Figure 3C**, **Supplementary Table 21**). We calculated the population attributable risk fraction and estimated that the proportion of COPD cases attributable to risk scores above the first GRS decile was 54.6% (95% CI: [50.6%, 58.4%]).

### Pleiotropy and phenome-wide association studies

As phenome-wide association studies (PheWAS) can provide evidence mimicking pharmacological interventions of drug targets in humans and informing drug development^42^, we undertook a PheWAS of 2,411 phenotypes in UK Biobank (**Online Methods, Figure 4**); 226 of the 279 sentinel variants were associated (FDR <1%) with one or more traits and diseases (excluding quantitative lung function traits). Eighty-five of the lung function signals were associated with standing height. In order to investigate whether the genetic association signals for lung function were driven by incomplete adjustment for height, we tested for correlation of effects on lung function in UK Biobank and height in the GIANT consortium for 247 of the 279 signals that had a proxy variant in GIANT^43^; there was no significant correlation (r=-0.096, **Supplementary Figure 5**). Additionally, the PheWAS revealed associations with body composition measures such as fat free mass (54 SNPs) and hip circumference (40 SNPs), as well as muscle strength (32 SNPs, grip strength). One hundred and fourteen of the 279 SNPs were associated with several quantitative measures of blood count, including eosinophil counts and percentages (25 SNPs). Twenty-five of our SNPs were also associated with asthma including 12 SNPs associated both with asthma and eosinophil measures. Five of these SNPs were in LD (r^2^>0.1) with a SNP reported for association both with asthma and eosinophil measures in previously published genome-wide association studies. To assess whether any of the lung function associations could be driven by an association with asthma, we compared the effect size estimated before and after exclusion of all self-reported asthma cases, observing remarkably similar estimates (**Supplementary Figure 6**) suggesting that the lung function associations we report are not primarily driven via known asthma signals.

**Figure 4:**
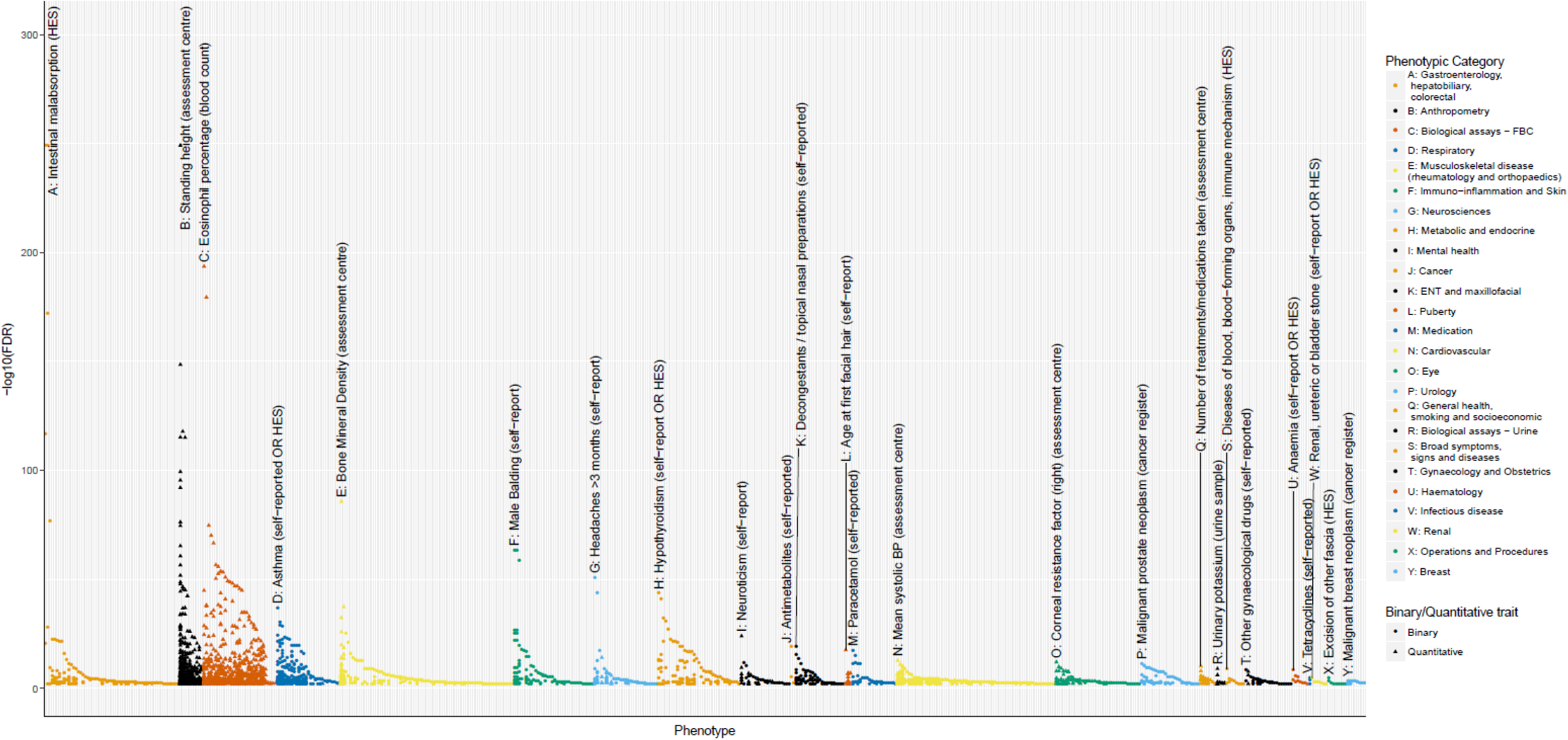
Individual PheWAS with 279 variants (traits passing FDR 1% threshold) Separate association of 279 variants with 2,411 traits (FDR<1%) in UK Biobank (n up to 379,337). In each category, the trait with the strongest association, i.e. highest −log_10_(FDR), is shown first, followed by other traits in that category in descending order of −log_10_(FDR). Categories are colour-coded, and outcomes are denoted with a circular or triangular point, according to whether they were coded as binary or quantitative. The top association per-category is labelled with its rsID number, and a plain English label describing the trait. The letter at the beginning of each label allows easy cross-reference with the categories labelled in the legend. Zoomed in versions of each category with visible trait names are available in **Supplementary Figure 9**.

We examined the specificity of genetic associations, given the potential for this to predict specificity of drug target modification, and found that 53 of the 279 signals were associated only with lung function and COPD-related traits. In contrast, three of our 279 signals were associated with over 100 traits across multiple categories – among these rs3844313, a known intergenic signal near *HLA-DQB1* was associated with 163 traits, and also had the strongest signal in the PheWAS, which was for association with intestinal malabsorption and coeliac disease.

In our 279-variant weighted GRS PheWAS analysis (**Supplementary Table 22**), we found association with respiratory traits including COPD, chronic bronchitis, emphysema, respiratory failure, corticosteroid use and both paediatric and adult-onset asthma (**Figure 5a**). The GRS was also associated with non-respiratory traits including coeliac disease, an intestinal autoimmune disorder (**Figure 5b**). These pleiotropic effects on risk of autoimmune diseases was further confirmed by analysis of previously reported GWAS (**Online Methods, Supplementary Table 23**) which showed overlapping single variant associations with Crohn’s disease, ulcerative colitis, psoriasis, systemic lupus erythematosus, IgA nephropathy, pediatric autoimmune disease and type 1 diabetes.

**Figure 5:**
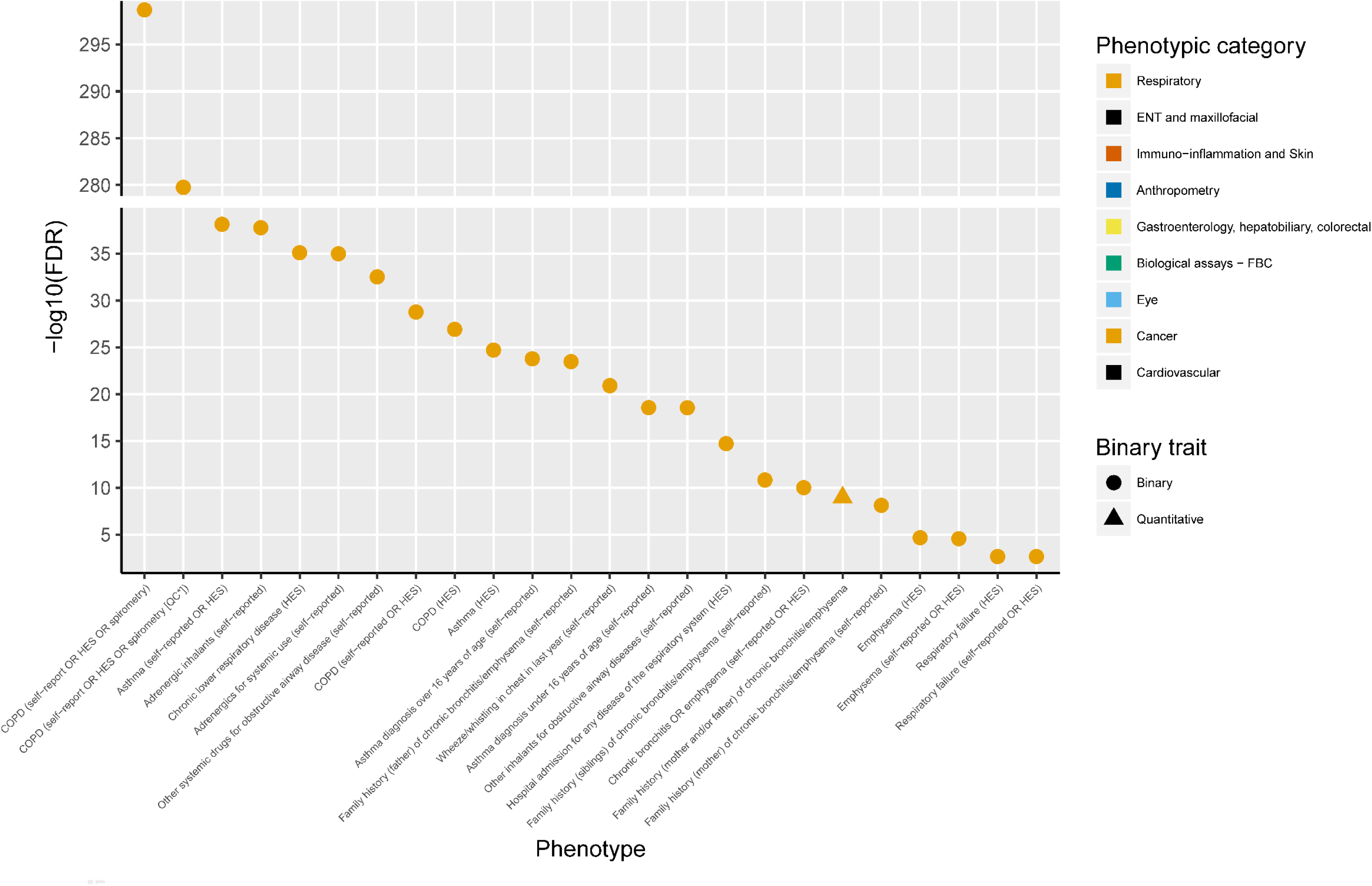
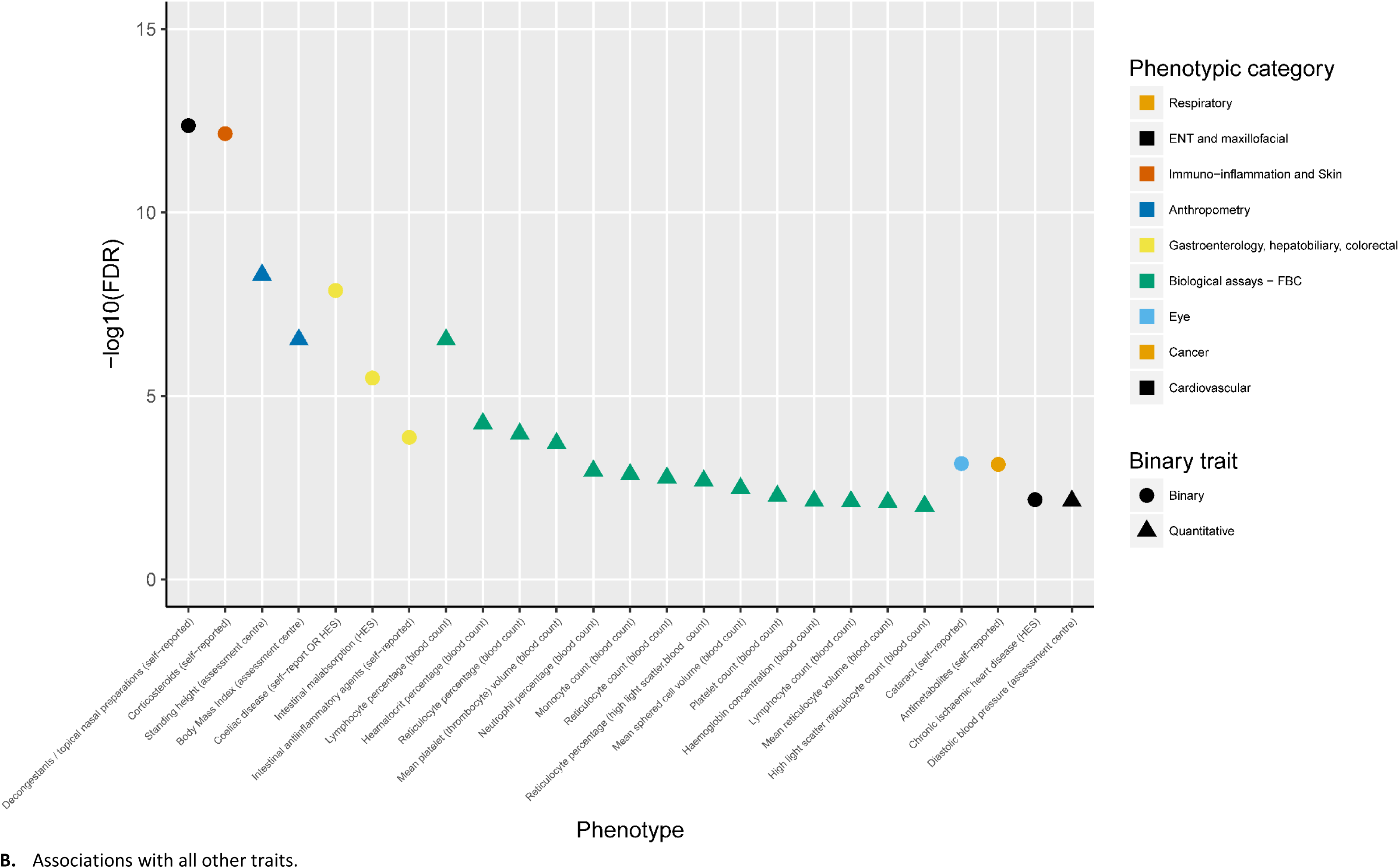
PheWAS with genetic risk score (traits passing FDR 1% threshold) Association of 279 variant weighted genetic risk score with 2,453 traits (FDR<1%) in UK Biobank (n up to 379,337). In each panel, the category with the strongest association, i.e. highest −log_10_(FDR), is shown first, followed by all other associations in that category, ordered by descending order of −log_10_(FDR). Sample sizes varied across traits and are available in **Supplementary Table 22**, along with the full summary statistics for each association, plus details of categorisation and plain English labels for each trait. Trait categories are colour coded, and outcomes are denoted with a circular or triangular point, according to whether they were coded as binary or quantitative. *QC refers to spirometry passing ERS/ATS criteria. **A.** Associations with respiratory traits.

## Discussion

The large sample size of our study, achieved by our refinement of the spirometry in UK Biobank and inclusion of the substantially expanded SpiroMeta consortium data set, has doubled the yield of lung function signals to 279. Fine-mapping of all new and previously reported signals, together with gene and protein expression analyses with improved tissue specificity and stringency, has implicated new genes and pathways, highlighting the importance of cilia development, TGFB-signalling via SMAD3, and elastic fibres in the aetiology of airflow obstruction. Many of the genes and pathways reported here contain druggable targets; we highlight examples where the genetic variants mimicking therapeutic modulation of targets may have opposing effects on lung function. We have developed and applied the first weighted GRS for lung function and tested it in deeply-phenotyped COPD case-control studies. Our GRS shows stronger association and larger effect size estimates (4.73 fold change in COPD risk between highest and lowest risk deciles) than a previous GRS in European ancestry populations^18^, as well as generalisability to African, South Asian and Chinese ancestry groups. We undertook the first comprehensive PheWAS for lung function signals, and report genetic variants with apparent specificity of effects and others with pleiotropic effects that might indicate shared biological pathways between different diseases.

For the first time in a GWAS of lung function, we report an enrichment of genes involved in ciliogenesis (including *KIAA0753*, *CDK2* and *CEP72*). Defects in primary cilia as a result of highly deleterious mutations in essential genes result in ciliopathies known to affect multiple organ systems. We found an enrichment of genes with a role in centriolar replication and duplication, core processes in primary and motile cilia formation. Mutations in *KIAA0753* cause the ciliopathies Joubert Syndrome and Orofaciodigital Syndrome^23^. Reduced airway motile cilia function impacting mucus clearance is a feature of COPD, but it has not been clear whether this is causal or the consequence of damage by external factors such as smoking or infection. Our findings suggest that impaired ciliary function might be a driver of the disease process. We have previously shown, through whole exome re-sequencing, an enrichment of rare variants in cilia-related genes in heavy smokers without airflow obstruction^44^.

New signals, implicating *ITGAV* and *GDF5*, as well as stronger support for *TGFB2* and *MFAP2* as likely causal genes, provide new genetic support for the importance of elastic fibre pathways in lung function and COPD^18^. The elastic fibres of the extracellular matrix are known to be disrupted in COPD^45^. As the breakdown of elastic fibres by neutrophil elastase leads to emphysema in individuals with alpha_1_-antitrypsin deficiency, we also assessed the association with the *SERPINA1* Z allele, which was not associated with lung function in our study (rs28929474, P=0.109 for FEV_1_/FVC in UK Biobank).

Smoking and genetic risk both have important effects on lung function and COPD. We found no interaction of smoking with individual lung function associated variants. Our weighted 279-SNP GRS showed no interaction with smoking status for FEV_1_/FVC, whilst a weak smoking-GRS interaction was observed for COPD susceptibility. Thus our findings are consistent with the effects of smoking and genetic risk being approximately additive on lung function (and multiplicative on COPD risk). Whilst the weighted 279-SNP GRS showed a strong association with COPD susceptibility, and a high attributable risk, we do not claim that this would represent an appropriate method of screening for COPD risk. Incorporation of the GRS into a risk model already comprising available clinical information (including age, sex, height and pack-years of smoking in COPDGene non-Hispanic Whites) leads to an increase in the area under the curve from 0.751 to 0.771, which although statistically significant (p=3.33×10^−10^) is of modest magnitude. Importantly, our findings demonstrate the high absolute risk among genetically susceptible smokers. Based on our estimated GRS relative risk and absolute risk estimates of COPD shown by Lokke *et al.*^46^, one would expect the highest GRS risk decile group of smokers to have an absolute risk of developing COPD by approximately 70 years of age of 82.4%, versus 17.4% for the lowest GRS decile.

The unprecedented sample size of UK Biobank as a single cohort has revolutionised genetic studies. We used two complementary study designs to maximise sample size for discovery and ensure robustness of findings by requiring independent support for association. Furthermore, through additional analysis of the spirometry data in UK Biobank and substantial expansion of the SpiroMeta consortium, we have markedly increased samples sizes to almost seven times those included in previous studies. As no lower MAF threshold was applied in our analyses, an overall threshold of P<5×10^−9^, as recommended for re-sequencing analyses of European ancestry individuals^47^, was applied. We identified the largest number of new signals in our more stringent two-stage design (“Tier 1”, 99 new signals). Amongst the signals that we report as “Tier 3” (and did not include in further analyses), all reached P<10^−3^ in UK Biobank and 183 met a less stringent threshold of P<0.05 in SpiroMeta.

Our study is the first to investigate genome-wide associations with PEF. PEF is determined by various physiological factors including lung volume, large airway calibre, elasticity of the lung and expiratory muscle strength, is used for monitoring asthma, and was incorporated in a recently evaluated clinical score for diagnosing COPD and predicting acute exacerbations of COPD^48^. Overall, 133 of the 279 signals were also associated with PEF (P<10^−5^) and for 15 signals (including 4 new signals), PEF was the most significantly associated trait. Of note, a signal near *SLC26A9*, a known cystic fibrosis modifier gene^49^, was highly significantly associated with PEF in UK Biobank (P=3.97×10^−66^) and was nominally significant in SpiroMeta (P=6.93×10^−3^), with consistent direction of effect, but did not meet the Tier 2 criteria (P<10^−3^ in each of SpiroMeta and UK Biobank). This could reflect the limited power for PEF in SpiroMeta (up to 24,218 for PEF compared to 79,055 for the other three traits).

Examining associations of a given genetic variant with a wide range of human phenotypes is a valuable tool in therapeutic target validation. As in our PheWAS, it can highlight variants which show associations with one or more respiratory traits that might be expected to demonstrate greater target specificity than variants associated with many traits. Additionally, in some instances, association with multiple traits may indicate the relevance of drug repurposing. Association of a given SNP with multiple traits does not necessarily imply shared aetiology, and further investigation is warranted. Our GRS PheWAS assesses broader genetic overlap between lung function and other traits and supports the evidence for some shared genetic determinants with autoimmune diseases.

In summary, our study has doubled the number of signals for lung function and, based on relating fine-mapped, annotated variants to gene and protein expression, epigenetic marks, gene sets, biological pathways and druggable proteins, it provides new understanding and resources of utility for the development of therapeutics. The 279-variant GRS we constructed was associated with a 4.71-fold increased relative risk of moderate-severe COPD between highest and lowest deciles, such that one would expect over 80% of smokers in the highest genetic risk decile to develop COPD. The GRS was also predictive of COPD across multiple ancestral groups. Our PheWAS highlights both expected and unexpected associations relevant to respiratory and other systemic diseases. Investigating the nature of the pleiotropic effects of some of these variants will be of benefit for drug target identification and validation.

## Online Methods

### Study Design Overview and rationale

For the two-stage approach, we firstly selected distinct signals of association (defined using conditional analyses) with one or more traits achieving P<5×10^−9^ in UK Biobank only (n up to 321,047). A threshold of P<5×10^−9^ was selected to maximise stringency of findings and to be consistent with currently recommended genome-wide significance thresholds for re-sequencing analyses of European ancestry individuals^50^. We then reported as new those signals which additionally met P<10^−3^ in SpiroMeta (N effective >70% of n up to 79,055; **Supplementary Note**, **Supplementary Figure 7**), with consistent directions of effect and term them “Tier 1” signals as they meet our highest level of stringency.

For the one-stage approach, we selected distinct signals of association (defined using conditional analyses) with one or more traits reaching P<5×10^−9^ in the meta-analysis of UK Biobank and SpiroMeta (n up to 400,102) and reported as new those which additionally met P<10^−3^ in both UK Biobank and SpiroMeta with a consistent direction of effect. We term these signals “Tier 2” as they meet our second-highest level of stringency.

All signals meeting either set of criteria described above, and that had not been previously published, were reported as new signals of association with lung function. Signals that reached P<5×10^−9^ in the meta-analysis of UK Biobank and SpiroMeta, had a consistent direction of effect in UK Biobank and SpiroMeta, but which did not reach P<10^−3^ in both UK Biobank and SpiroMeta are presented as “Tier 3” and were not included in further analyses.

### UK Biobank

The UK Biobank data resource is described elsewhere (see URLs). Individuals were selected for inclusion in this study if they met the following criteria: (i) had complete data for age, sex, height and smoking status; (ii) had spirometry meeting quality control requirements (based on analyses of acceptability, reproducibility and blow curve metrics; **Supplementary Note**); (iii) had genome-wide imputed genetic data and; (iv) were of European ancestry based on genetic data (**Supplementary Note**; **Supplementary Figure 1**). Genotyping was undertaken using the Affymetrix Axiom^®^ UK BiLEVE and UK Biobank arrays^13^. Genotypes were imputed to the Haplotype Reference Consortium panel^51^ (**Supplementary Note**), and retained if minor allele count ≥3 and imputation quality (info) > 0.5. A total of 321,047 individuals were included in this analysis (**Supplementary Table 1**).

Residuals from linear regression of each trait (FEV_1_, FVC, FEV_1_/FVC and PEF) against age, age^2^, sex, height, smoking status (ever/never) and genotyping array were ranked and inverse-normal transformed to obtain adjusted, normally distributed Z-scores. These Z-scores were then used for genome-wide association testing under an additive genetic model using BOLT-LMM v2.3^20^. Principal components were not included as BOLT-LMM uses a linear mixed model to account for relatedness and fine-scale population structure.

Linkage disequilibrium (LD) score regression implemented in LDSC^52^ was used to estimate inflation of test statistics due to confounding. Genomic control was applied, adjusting all test statistics by LD score regression intercepts: 1.12 for FEV_1_, 1.14 for FVC, 1.19 for FEV_1_/FVC and 1.13 for PEF (**Supplementary Figure 8**; **Supplementary Table 24**).

### SpiroMeta consortium

The SpiroMeta consortium meta-analysis was comprised of a total of 79,055 individuals from 22 studies. Thirteen studies (n=21,436 individuals) were imputed to the 1000 Genomes Project Phase 1 reference panel^53^ (B58C [T1DGC and WTCCC], BHS1&2, three Croatian studies [CROATIA-Korcula, CROATIA-Split and CROATIA-Vis], Health 2000, KORA F4, KORA S3, LBC1936, NSPHS, ORCADES, SAPALDIA and YFS and 9 studies (n=61,682 individuals) were imputed to the Haplotype Reference Consortium (HRC) panel^54^ (EPIC [obese cases and population-based studies], GS:SFHS, NFBC1966, NFBC1986, PIVUS, SHIP, SHIP-TREND, UKHLS and VIKING). See **Supplementary Tables 2** and **3** for the definitions of all abbreviations, study characteristics, details of genotyping platforms and imputation panels and methods). Measurements of spirometry for each study are described in the **Supplementary Note**.

In each study, linear regression models were fitted for each lung function trait (FEV_1_, FEV_1_/FVC, FVC and PEF, where available), with adjustment for age, age^2^, sex and height. For studies with unrelated individuals, these models were fitted separately in ever smokers and never smokers, with additional adjustment for principal components of ancestry. Studies with related individuals fitted mixed models in all individuals to account for relatedness, with ever smoking status as a covariate.

In all studies, rank-based inverse normal transformations were undertaken on the residuals, with these transformed residuals used as the phenotype for association testing under an additive genetic model (**Supplementary Table 3**).

In the study level results, variants were excluded if they had a very low MAC (**Supplementary Table 3**) or imputation quality (info) <0.3. In studies with unrelated individuals, the ever and never smokers results were combined, using inverse variance weighted meta-analysis, to give an overall study result. Genomic control was then applied to all study level results, before combining results across all studies using inverse variance weighted meta-analysis. LD score regression intercepts for the meta-analysis were close to 1 (**Supplementary Figure 8**; **Supplementary Table 24**) and so genomic control was not applied.

### Meta-analyses

A total of 19,871,028 variants (imputed or genotyped) in both UK Biobank and SpiroMeta were meta-analysed using inverse-variance weighted fixed effect meta-analysis, and no further genomic control was applied as LD score regression intercepts were close to 1 (**Supplementary Table 24**).

### Selection of new signals using conditional analyses

All SNPs ±1Mb were extracted around each sentinel variant. GCTA^55^ was then used to perform stepwise conditional analysis to select independently associated SNPs within each 2Mb region. Any secondary signals identified within each 2Mb region were required to meet Tier 1 or Tier 2 criteria (described above) after conditioning on the primary sentinel variant. A combined list of distinct lung function signals was then made across the 4 phenotypes, FEV_1_, FVC, FEV_1_/FVC and PEF as follows: where sentinel variants for 2 signals for different phenotypes were in high LD (r^2^ > 0.5), we retained the most significant variant; where 2 signals were in moderate LD (0.1 > r^2^ > 0.5), we retained variants if, after conditional analysis, they still met the Tier 1 or Tier 2 threshold; for signals in low LD (r^2^ < 0.1) we retained both variants. We then used the same criteria to identify a subset of new signals which were distinct from previously published independent signals (see below).

### Assessment of previously reported lung function signals

We identified 184 autosomal signals from previous GWAS analyses of lung function and COPD^1,4–14^. After LD pruning (keeping only those signals with LD of r^2^ < 0.1), we removed 24 non-independent SNPs, leaving 160 previously reported independent signals. Of 6 previously reported signals in the HLA region, we included only the 3 independent lung function HLA signals reported from conditional analysis using all imputed HLA genotypes^18^: *AGER* (rs2070600), *HLA-DQB1* (rs114544105) and near *ZNF184* (rs34864796) leaving 157 signals.

We confirmed association of previously reported signals in our data if they met any of three criteria: (i) the previously reported sentinel was associated (P<10^−5^) with any lung function trait in UK Biobank; (ii) a proxy for the previously reported sentinel with r^2^>0.5 was associated (P<10^−5^) with any lung function trait in UK Biobank; (iii) a proxy for the previously reported sentinel with r^2^>0.1 was associated with any lung function trait meeting tier 1 or tier 2 criteria (**Supplementary Figure 3**).

### Effect on COPD susceptibility – genetic risk score in multiple ancestries

To test association of all lung function signals and COPD susceptibility, we constructed a 279-variant weighted GRS comprising the 139 novel and 140 previously reported signals; we used the previously reported sentinel SNP for published signals. Weights were derived using the FEV_1_/FVC ratio decreasing (i.e. COPD risk *increasing*) alleles. For previously reported signals (n=140), results from the UK Biobank analysis were used to derive weights for the 94 signals that were not discovered using UK Biobank data and weights were taken from SpiroMeta for 46 signals where UK Biobank was included in the discovery of those signals. For novel signals identified in this study, weights were taken from SpiroMeta for two-stage (tier 1) signals (n=99), and the smallest absolute effect size from either of UK Biobank or SpiroMeta was used for one-stage (tier 2) signals (n=40) (**Supplementary Table 25**). For the weighted GRS the number of risk alleles at each variant was multiplied by its weight.

The GRS was first calculated in unrelated individuals (KING kinship coefficient of < 0.0884) within 6 ancestral groups of UK Biobank: Europeans, South Asians, Africans, Chinese, Mixed African and Europeans, and Mixed Other (total sample of unrelated individuals across six ancestries: 323,001) using PLINK. Weights and alleles were as described above. COPD was defined as FEV_1_/FVC < 0.7 and FEV_1_ < 0.8 of the predicted value, i.e. GOLD stage 2-4 categorisation. Associations with the GRS were then tested using COPD (in ancestral groups with at least 100 COPD cases) and FEV_1_/FVC as the outcomes.

In addition, we calculated the GRS in individuals from the China Kadoorie Biobank (CKB). Four of the 279 SNPs were not available in CKB (rs1800888, rs56196860, rs72724130 and rs77672322), and for 12 SNPs, proxies were used (minimum r^2^=0.3). Analyses were undertaken in all COPD GOLD stage 2-4 cases (FEV_1_/FVC < 0.7 and FEV_1_ < 0.8 of the predicted value, in 6,013 cases and 69,567 controls), against an unbiased set of population controls. The GRS was also tested for association with FEV_1_/FVC in CKB (n=72,796).

Logistic regression of COPD case-control status with the GRS in UK Biobank and China Kadoorie Biobank assumed an additive genetic effect and was adjusted for age, age^2^, sex, height, and smoking (**Supplementary Table 18**). Ten principal components were also included in UK Biobank analyses. In China Kadoorie Biobank, analyses were stratified by geographical regions and then meta-analysed using an inverse-variance fixed effect model. Linear models assessing the association with FEV_1_/FVC were fitted using the same transformed outcome as in the main GWAS analysis.

We then tested association in 5 European ancestry COPD case-control studies: COPDGene (Non-Hispanic White Population) (3,068 cases and 2,110 controls), ECLIPSE (1,713 cases and 147 controls), GenKOLS (836 cases and 692 controls), NETT-NAS (374 cases and 429 controls) and SPIROMICS (988 cases and 537 controls) (**Supplementary Table 19**). In addition, we tested this GRS in the COPDGene African American population study (910 cases and 1,556 controls). Logistic regression models using COPD as outcome and the GRS as exposure were adjusted for age, age^2^, sex, height, and principal components (**Supplementary Table 20**).

Next, we divided individuals in the external COPD case-control studies into deciles according to their values of the weighted GRS. This was undertaken separately by study group, and for each decile logistic models were fitted, comparing the risk of COPD for members of each decile group compared to those in the lowest decile (i.e. those with lowest values of the weighted GRS). Covariates were as for the COPD analyses. Results were combined across European-ancestry study groups by fixed effect meta-analysis (**Supplementary Table 21**).

We calculated the population attributable risk fraction (PARF) as follows:

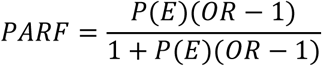

where *P(E)* is set to 0.9, i.e. the probability of carrying more risk alleles than those in the lowest risk score decile of the risk score (the ‘probability of the exposure’). *OR* refers to the odds of having COPD in individuals across deciles 2 to 10 of the risk score compared to the odds of having COPD for individuals in the lowest decile (decile 1) of the risk score (**Supplementary Note**).

### Effects on smoking behaviour

As our discovery GWAS in UK Biobank was adjusted for ever vs. never smoking status, and not for pack years of smoking (pack years information was missing for 32% of smokers), we evaluated whether any signals of association with lung function might be driven by an association with smoking behaviour by testing for association with smoking initiation (123,890 ever smokers vs. 151,706 never smokers) and cigarettes per day (n=80,015) in UK Biobank (full methods in **Supplementary Note**). We also tested for association with lung function in never smokers only (n=173,658). We excluded any signals associated with smoking behaviour (**Supplementary Table 6**), but not with lung function in never smokers.

### Smoking interaction

For associated variants (new and previously reported), we repeated association testing for lung function separately in UK Biobank and SpiroMeta (up to 176,701 ever smokers and 197,999 never smokers), and tested for an interaction effect with smoking using the Welch test (**Supplementary Note**). A threshold of P<1.79×10^−4^ (Bonferroni corrected for 279 tests) indicated significance.

We further tested for interaction between the weighted GRS and smoking, within 303,619 unrelated individuals of European ancestry in UK Biobank, using COPD and FEV_1_/FVC as outcomes (the FEV_1_/FVC phenotype was pre-adjusted for age, age^2^, sex, and height, and the residuals transformed as per the main GWAS analysis). For COPD (defined as FEV_1_/FVC<0.7, and FEV_1_ <80% predicted) the following logistic model was fitted:

> *COPD ̃ genotyping array + 10 principal components + age + age^2^ + sex + height + smoking status + weighted risk score +* (*smoking status* × *weighted risk score*).

For FEV_1_/FVC the following linear model was fitted:

> *FEV_1_/FVC ^∼^ genotyping array + 10 principal components + smoking status + weighted risk score + (smoking status x weighted risk score).*

### Proportion of variance explained

We calculated the proportion of variance explained by each of the previously reported (n=140) and new variants (n=139) associated with lung function using the formula:

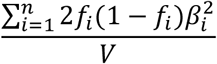

where n is the number of variants *f_i_* and *β_i_* are the frequency and effect estimate of the i’th variant, and V is the phenotypic variance (always 1 as our phenotypes were inverse-normal transformed). We used the same unbiased effect estimates (β) as used to calculate GRS weights at the same set of 279 sentinel variants used for the GRS, which uses either UK Biobank or SpiroMeta effect estimates (described above). Our previously published estimate of proportion of variance explained^18^ used effect estimates derived from UK Biobank. We assumed a heritability of 40%^56,57^ to estimate the proportion of additive polygenic variance.

### Fine-mapping

A Bayesian method^58^ was used to fine-map lung-function-associated signals to the set of variants that were 99% likely to contain the underlying causal variant (assuming that the causal variant has been analysed). This was undertaken for new signals and for previously reported signals reaching P<10^−5^ in UK Biobank. For the previously reported signals, the top sentinel variant from the current analysis in UK Biobank was used, instead of the previously reported variant. We used a value of 0.04 for the prior W in the approximate Bayes factor formula^59^. Effect sizes and standard errors for fine-mapping were obtained from an inverse variance weighted meta-analysis of UK Biobank and SpiroMeta (n up to 400,102). Signals in the HLA region were not included.

### Implication of potentially causal genes

#### Annotation of deleterious variants

Variants in the 99% credible sets were checked for predicted functional effect if they were annotated as “exonic”, “splicing”, “ncRNA_exonic”, “5’ UTR” or “3’ UTR” (untranslated region) by ANNOVAR^60^. We then used SIFT, PolyPhen-2 (implemented using the Ensembl GRCh37 Variant Effect Predictor, see URLs, accessed 1 February 2018) and FATHMM^61^ to annotate missense variants, and CADD (also implemented using VEP) to annotate non-coding variation. Variants were annotated as deleterious in our study if they were labelled ‘deleterious’ by SIFT, ‘probably damaging’ or ‘possibly damaging’ by PolyPhen-2, ‘damaging’ by FATHMM (specifying the ‘Inherited disease’ option of the coding variants methods, and setting the prediction algorithm to ‘Unweighted’) or had a CADD scaled score ≥20 ^4^. The union of the four methods was taken to establish the number of potentially deleterious variants and their unique genes.

#### Gene expression and protein levels

At each novel and previously reported signal, the sentinel variant and 99% credible set^58^ were used to query three eQTL resources: lung eQTL (n=1,111)^13^, blood eQTL (n=4,896)^62^ and GTEx (V7; with n up to 388 depending on tissue: Artery Aorta (n=267), Artery Coronary (n=152), Artery Tibial (n=388), Colon Sigmoid (n=203), Colon Transverse (n=246), Esophagus Gastroesophageal Junction (n=213), Esophagus Muscularis (n=335), Lung (n=383), Small Intestine Terminal Ileum (n=122), Stomach (n=237), and Whole Blood (n=369))^63^, and one blood pQTL resource (n=3,301)^34^.

A gene was classified as a ‘putative causal gene’ if the sentinel SNP or any SNP in the respective 99% credible set was associated with expression of this gene or its protein levels (FDR<5% for eQTL, P<5.03×10^−8^ [for 276 tests at 3,600 proteins] for pQTL) and if the GWAS sentinel SNP or any SNP in the respective 99% credible set was also the variant most strongly associated with expression of the respective gene or level of the respective protein (i.e. the sentinel eQTL/pQTL SNP) in one or more of the eQTL and pQTL data sets. The HLA region was excluded from these analyses.

### Pathway analysis

We tested for enrichment of genes identified via variant function annotation, gene expression or protein level analyses in pathway and gene set ontology databases using ConsensusPathdb. Pathways or gene sets represented entirely by genes implicated by the same association signal were excluded. Gene sets and pathways with FDR<5% are reported.

### Functional enrichment analyses

We tested for cell-specific enrichment of lung function associated variants in regulatory regions using FORGE^35^ (v1.1). One thousand background SNP set repetitions were used. Thresholds P<1.68×10^−4^ (FDR<2%; >99^th^ percentile) and P<3.37 × 10^−5^ (FDR<0.5%; >99.9^th^ percentile) were taken as being ‘indicative’ and ‘significant’, respectively. FORGE analysis was carried out for the cell lines in the RoadMap Epigenome project^33^ (n=299 cell lines) and ENCODE projects^64^ (n=125) separately.

Using DeepSEA^36^, we analysed all SNPs in the 99% credible set for predicted chromatin effects. We reported effects for any chromatin effect and lung-related cell line that had an E-value<0.05 (i.e. the expected proportion of SNPs with a larger predicted effect based on empirical distributions of predicted effects for 1000 Genomes SNPs) and an absolute difference in probability of >0.1 (threshold for “high confidence”) between the reference and alternative allele.

### Drug targets

Genes identified as potentially causal using eQTL, pQTL or variant annotation were interrogated against the gene-drug interactions table of the Drug-Gene Interactions Database (DGIDB) (see URLs), accessed 16^th^ October 2017. Drugs were mapped to CHEMBL IDs (see URLs), and indications (as MeSH headings) were added.

### Phenome-wide association studies

To identify whether any of the new or previously reported signals overlap with signals of association for other traits and diseases, the 279 variant weighted GRS was calculated in UK Biobank samples (n up to 379,337) and a phenome-wide association study (PheWAS) across all available traits was performed, with the risk score as the exposure. Traits included UK Biobank baseline measures (from both questionnaires and physical measures), self-reported medication usage, and operative procedures, as well as those captured in Office of Population Censuses and Surveys codes from the electronic health record. We also included self-reported disease variables and those from hospital episode statistics (ICD-10 codes truncated to three-character codes and combined in block and chapter groups) as well as combining both self-report and hospital diagnosed diseases where possible to maximise power. The GRS analysis included 2,453 traits, of which 2,411 were also included in the single-variant analysis (traits with >200 cases were included for the individual SNP PheWAS, whereas traits with >50 cases were included in the risk score PheWAS). Analyses were conducted in unrelated European ancestry individuals (KING kinship coefficient of <0.0442), and were adjusted for age, sex, genotyping array, and ten principal components. Logistic models were fitted for binary outcome, and linear models were fitted for quantitative outcomes. False discovery rates were calculated according to the number of the traits in each analysis (2,453 or 2,411, for the risk score and single-variant PheWAS, respectively).

In addition, the sentinel variants and variants within the 99% credible sets were queried against the GWAS catalog^65^ (see URLs, accessed 5th February 2018) and GRASP^66^ (see URLs, accessed 6th February 2018) for reported associations significant at P<5×10^−8^. Associations relating to methylation, expression, metabolite or protein levels, as well as lung function and COPD, were not included.

### Data availability statement

UK Biobank GWAS summary statistics will be available via UK Biobank (http://www.ukbiobank.ac.uk/). SpiroMeta GWAS summary statistics, and single-variant PheWAS results will be made available by request.

## URLs

UK Biobank: http://www.ukbiobank.ac.uk

Variant Effect Predictor: https://www.ensembl.org/vep

Drug-Gene Interactions Database (DGIDB): http://www.dgidb.org/data/

CHEMBL: https://www.ebi.ac.uk/chembl/drug/indications

GWAS catalog: https://www.ebi.ac.uk/gwas/

GRASP: https://grasp.nhlbi.nih.gov/Overview.aspx

## Study contributions

All authors critically reviewed the manuscript prior to submission.

Contributed to the conception and design of the study: K.S., U.S.S.G., S.K., S.M.K., T.L., P.S.B., T.H.B., E.R.B., Y.B., Z.C., J.D.C., J.D., D.L.D., C.G., A.G., K.H., J.D.H., J.Hokanson, P.J., C.L., L.Li, N.L., J.C.M., H.R., I.Sayers, D.D.S., R. T-S., J.C.W., P.W., L.M.Y., O.T.R., M.K., O.P., U.G., I.R., I.J.D., N.M.P., H.S., A.L.J., J.F.W., E.Z., M.J., N.W., A.S.B., R.A.S., D.A.M., M.H.C., D.P.S., I.P.H., M.D.T., L.V.W.

Undertook data analysis: N.S., A.L.G., A.M.E., V.E.J., B.D.H., C.M., C.Batini, K.A.F., K.S., P.S., Xingnan Li, R.B., N.F.R., M.O., J.Zhao, M.W., S.W., K.A.K., J.P.C., B.B.S., J.Zhou, J.Hui, M.I., S.E.H., J.M., S.E., I.Surakka, V.V., T.L., R.J.A., C.Brandsma, F.D., J.D.H., P.K.J., Xuan Li, A.Mahajan, J.C.M., D.C.N., M.M.P., D.Prokopenko, D.Q., R.R., H.R., D.S., P.R.H.J.T., M.V., L.M.Y., O.G.T., N.M.P., N.W., E.K.S., C.H., A.P.M., A.S.B., R.A.S., M.H.C., D.P.S., M.D.T., L.V.W.

Contributed to data acquisition and/or interpretation: N.S., A.L.G., A.M.E., V.E.J., C.M., C.Batini, K.A.F., K.S., P.S., Xingnan Li, N.F.R., M.O., M.W., K.A.K., B.B.S., S.K., M.I., R.J.A., J.D., F.D., R.E., C.G., A.G., A.L.H., J.D.H., G.H., P.K.J., C.L., Xuan Li, K.L., L.Lind, J.L., J.C.M., A.Murray, R.P., M.M.P., M.L.P., D.Porteous, D.Prokopenko, D.Q., R.R., H.R., I.Sayers, B.H.S., M.S., L.M.Y., O.G.T., N.M.P., H.S., J.F.W., B.S., M.J., N.W., C.H., A.P.M., A.S.B., R.A.S., R.G.W., M.H.C., D.P.S., I.P.H., M.D.T., L.V.W. Drafted the manuscript: N.S., A.L.G., A.M.E., I.P.H., M.D.T., L.V.W.

## Funding and Acknowledgments

### The following authors report specific personal funding from the following grants

B.D.H.: NIH K08 HL136928, Parker B. Francis Research Opportunity Award

M.W.: EU Horizon2020 (633212 ALEC)

U.S.S.G.: Economic and Social Research Council (ES/H029745/1)

B.B.S.: Cambridge School of Clinical Medicine MB-PhD programme and MRC/Sackler Prize PhD Studentship (MR/K50127X/1).

J.D.: J.D. is a BHF Professor, European Research Council Senior Investigator, and NIHR Senior Investigator.

I.Sayers: MRC (G1000861)

D.S.: D.S. is supported by a VA Research Career Scientist award. E.Z.: Wellcome Trust (WT098051)

A.P.M.: Wellcome Trust (WT098017 & WT064890)

M.H.C.: This work was supported by NHLBI grants R01HL113264, R01HL137927 (M.H.C. and E.K.S.), R01HL135142 (M.H.C.). The content is solely the responsibility of the authors and does not necessarily represent the official views of the NIH. The funding body has no role in the design of the study and collection, analysis, and interpretation of data and in writing the manuscript.

I.P.H.: The research was partially supported by the NIHR Nottingham Biomedical Research Centre; the views expressed are those of the author(s) and not necessarily those of the NHS, the NIHR or the Department of Health.

M.D.T.: M.D. Tobin is supported by a Wellcome Trust Investigator Award (WT202849/Z/16/Z). M.D. Tobin and L.V. Wain have been supported by the MRC (MR/N011317/1). The research was partially supported by the NIHR Leicester Biomedical Research Centre; the views expressed are those of the author(s) and not necessarily those of the NHS, the NIHR or the Department of Health.

L.V.W.: L.V. Wain holds a GSK/British Lung Foundation Chair in Respiratory Research.

### Funding and acknowledgment statements from individual cohorts

China Kadoorie Biobank: Kadoorie Charitable Foundation in Hong Kong; the UK Wellcome Trust (grant numbers 202922/Z/16/Z, 088158/Z/09/Z, 104085/Z/14/Z); Chinese Ministry of Science and Technology (grant number 2011BAI09B01); National Natural Science Foundation of China (Grant numbers 81390540, 81390541, 81390544); UK Medical Research Council (grant numbers MC_PC_13049, MC_PC_14135); GlaxoSmithKline; BHF Centre of Research Excellence, Oxford (grant number RE/13/1/30181); British Heart Foundation; and Cancer Research UK. The China Kadoorie Biobank participants; CKB project staff based at Beijing, Oxford and the 10 regional centres; the China National Centre for Disease Control and Prevention (CDC) and its regional offices for assisting with the fieldwork. We acknowledge the assistance of BGI (Shenzhen, China) for conducting DNA extraction and genotyping.

COPDGene: See Supplementary Note.

ECLIPSE: The ECLIPSE study (NCT00292552; GSK code SCO104960) was funded by GSK.

GenKOLS: The Norway GenKOLS study (Genetics of Chronic Obstructive Lung Disease, GSK code RES11080) was funded by GSK.

NETTNAS: The National Emphysema Treatment Trial was supported by the NHLBI N01HR76101, N01HR76102, N01HR76103, N01HR76104, N01HR76105, N01HR76106, N01HR76107, N01HR76108, N01HR76109, N01HR76110, N01HR76111, N01HR76112, N01HR76113, N01HR76114, N01HR76115, N01HR76116, N01HR76118 and N01HR76119, the Centers for Medicare and Medicaid Services and the Agency for Healthcare Research and Quality. The Normative Aging Study is supported by the Cooperative Studies Program/ERIC of the US Department of Veterans Affairs and is a component of the Massachusetts Veterans Epidemiology Research and Information Center (MAVERIC).

SPIROMICS: NIH/NHLBI grants: HHSN268200900013C, HHSN268200900014C, HHSN268200900015C, HHSN268200900016C, HHSN268200900017C, HHSN268200900018C, HHSN268200900019C (Eugene Bleecker), HHSN268200900020C. Supplemented by contributions made through the Foundation for the NIH from AstraZeneca; Bellerophon Pharmaceuticals; Boehringer-Ingelheim Pharmaceuticals, Inc; Chiesi Farmaceutici SpA; Forest Research Institute, Inc; GSK; Grifols Therapeutics, Inc; Ikaria, Inc; Nycomed GmbH; Takeda Pharmaceutical Company; Novartis Pharmaceuticals Corporation; Regeneron Pharmaceuticals, Inc; and Sanofi. The authors thank the SPIROMICS participants and participating physicians, investigators and staff for making this research possible. More information about the study and how to access SPIROMICS data is at www.spiromics.org. We would like to acknowledge the following current and former investigators of the SPIROMICS sites and reading centers: Neil E Alexis, PhD; Wayne H Anderson, PhD; R Graham Barr, MD, DrPH; Eugene R Bleecker, MD; Richard C Boucher, MD; Russell P Bowler, MD, PhD; Elizabeth E Carretta, MPH; Stephanie A Christenson, MD; Alejandro P Comellas, MD; Christopher B Cooper, MD, PhD; David J Couper, PhD; Gerard J Criner, MD; Ronald G Crystal, MD; Jeffrey L Curtis, MD; Claire M Doerschuk, MD; Mark T Dransfield, MD; Christine M Freeman, PhD; MeiLan K Han, MD, MS; Nadia N Hansel, MD, MPH; Annette T Hastie, PhD; Eric A Hoffman, PhD; Robert J Kaner, MD; Richard E Kanner, MD; Eric C Kleerup, MD; Jerry A Krishnan, MD, PhD; Lisa M LaVange, PhD; Stephen C Lazarus, MD; Fernando J Martinez, MD, MS; Deborah A Meyers, PhD; John D Newell Jr, MD; Elizabeth C Oelsner, MD, MPH; Wanda K O’Neal, PhD; Robert Paine, III, MD; Nirupama Putcha, MD, MHS; Stephen I. Rennard, MD; Donald P Tashkin, MD; Mary Beth Scholand, MD; J Michael Wells, MD; Robert A Wise, MD; and Prescott G Woodruff, MD, MPH. The project officers from the Lung Division of the National Heart, Lung, and Blood Institute were Lisa Postow, PhD, and Thomas Croxton, PhD, MD.

DeepSEA: The DeepSEA project was primarily supported by US National Institutes of Health (NIH) grants R01 GM071966 to O.G.T and supported in part by US NIH grant P50 GM071508. O.G.T. is a senior fellow of the Genetic Networks program of the Canadian Institute for Advanced Research (CIFAR). Jian Zhou and Olga G Troyanskaya carried out SNV effect prediction analyses in an updated version of DeepSEA (deepsea.princeton.edu), which contained more lung-related cell lines than the original version published in Nature Methods in August 2015 (www.nature.com/articles/nmeth.3547).

GSK: This research has been conducted using the UK Biobank Resource under application number 26041.

B58C: Genotyping for the B58C-WTCCC subset was funded by the Wellcome Trust (076113/B/04/Z). The B58C-T1DGC genotyping utilized resources provided by the Type 1 Diabetes Genetics Consortium, a collaborative clinical study sponsored by the National Institute of Diabetes and Digestive and Kidney Diseases (NIDDK), National Institute of Allergy and Infectious Diseases (NIAID), National Human Genome Research Institute (NHGRI), National Institute of Child Health and Human Development (NICHD), and Juvenile Diabetes Research Foundation International (JDRF) and supported by U01 DK062418. B58C-T1DGC GWAS data were deposited by the Diabetes and Inflammation Laboratory, Cambridge Institute for Medical Research (CIMR), University of Cambridge, which is funded by Juvenile Diabetes Research Foundation International, the Wellcome Trust and the National Institute for Health Research Cambridge Biomedical Research Centre; the CIMR is in receipt of a Wellcome Trust Strategic Award (079895). The B58C-GABRIEL genotyping was supported by a contract from the European Commission Framework Programme 6 (018996) and grants from the French Ministry of Research.

BHS: Generous support for the 1994/5 follow-up study from Healthway, Western Australia, and support from The Great Wine Estates of the Margaret River region of Western Australia. The study also acknowledges the numerous Busselton community volunteers who assisted with data collection and the study participants from the Shire of Busselton.

CROATIA-Korcula/Split/Vis: MRC, the Ministry of Science, Education and Sport in the Republic of Croatia (216-1080315-0302) and the Croatian Science Foundation (grant 8875) (funding to I. Rudan, C. Hayward, S.M. Kerr, O. Polasek, V. Vitart, and J. Marten).

H2000: Medical Research Fund of the Tampere University Hospital.

KORA F4 and KORA S3: The KORA study was initiated and financed by the Helmholtz Zentrum München – German Research Center for Environmental Health, which is funded by the German Federal Ministry of Education and Research (BMBF) and by the State of Bavaria. Furthermore, KORA research was supported within the Munich Center of Health Sciences (MC-Health), LudwigMaximilians-Universität, as part of LMUinnovativ and by the Competence Network Asthma and COPD (ASCONET), network COSYCONET (subproject 2, BMBF FKZ 01GI0882) funded by the German Federal Ministry of Education and Research (BMBF).

LBC1936: Phenotype collection in the Lothian Birth Cohort 1936 was supported by Age UK (The Disconnected Mind project). Genotyping was funded by the Biotechnology and Biological Sciences Research Council (BBSRC). The work was undertaken by The University of Edinburgh Centre for Cognitive Ageing and Cognitive Epidemiology, part of the cross council Lifelong Health and Wellbeing Initiative (MR/K026992/1). Funding from the BBSRC and MRC is gratefully acknowledged.

NSPHS: Swedish Medical Research Council (K2007-66X-20270-01-3, 2011-5252, 2012-2884 and 2011-2354), the Foundation for Strategic Research (SSF). NSPHS as part of European Special Populations Research Network (EUROSPAN) was also supported by the European Commission FP6 STRP (01947, LSHG-CT-2006-01947).

ORCADES: Supported by the Chief Scientist Office of the Scottish Government (CZB/4/276, CZB/4/710), the Royal Society, the MRC Human Genetics Unit, Arthritis Research UK and the European Union framework program 6 EUROSPAN project (contract no. LSHG-CT-2006-018947). ORCADES DNA extractions were performed at the Wellcome Trust Clinical Research Facility in Edinburgh.

SAPALDIA: Swiss National Science Foundation (33CS30-148470/1&2, 33CSCO-134276/1, 33CSCO-108796, 324730_135673, 3247BO-104283, 3247BO-104288, 3247BO-104284, 3247-065896, 3100- 059302, 3200-052720, 3200-042532, 4026-028099, PMPDP3_129021/1, PMPDP3_141671/1), the Federal Office for the Environment, the Federal Office of Public Health, the Federal Office of Roads and Transport, the canton’s government of Aargau, Basel-Stadt, Basel-Land, Geneva, Luzern, Ticino, Valais, and Zürich, the Swiss Lung League, the canton’s Lung League of Basel Stadt/ Basel Landschaft, Geneva, Ticino, Valais, Graubünden and Zurich, Stiftung ehemals Bündner Heilstätten, SUVA, Freiwillige Akademische Gesellschaft, UBS Wealth Foundation, Talecris Biotherapeutics GmbH, Abbott Diagnostics, European Commission 018996 (GABRIEL) and the Wellcome Trust (WT 084703MA). The SAPALDIA study could not have been done without the help of the study participants, technical and administrative support and the medical teams and field workers at the local study sites. Local fieldworkers: Aarau: S Brun, G Giger, M Sperisen, M Stahel, Basel: C Bürli, C Dahler, N Oertli, I Harreh, F Karrer, G Novicic, N Wyttenbacher, Davos: A Saner, P Senn, R Winzeler, Geneva: F Bonfils, B Blicharz, C Landolt, J Rochat, Lugano: S Boccia, E Gehrig, MT Mandia, G Solari, B Viscardi, Montana: AP Bieri, C Darioly, M Maire, Payerne: F Ding, P Danieli A Vonnez, Wald: D Bodmer, E Hochstrasser, R Kunz, C Meier, J Rakic, U Schafroth, A Walder.

YFS: The Young Finns Study has been financially supported by the Academy of Finland: grants 286284, 134309 (Eye), 126925, 121584, 124282, 129378 (Salve), 117787 (Gendi), and 41071 (Skidi); the Social Insurance Institution of Finland; Competitive State Research Financing of the Expert Responsibility area of Kuopio, Tampere and Turku University Hospitals (grant X51001); Juho Vainio Foundation; Paavo Nurmi Foundation; Finnish Foundation for Cardiovascular Research; Finnish Cultural Foundation; Tampere Tuberculosis Foundation; Emil Aaltonen Foundation; Yrjö Jahnsson Foundation; Signe and Ane Gyllenberg Foundation; Diabetes Research Foundation of Finnish Diabetes Association; and EU Horizon 2020 (grant 755320 for TAXINOMISIS).

EPIC: Cancer Research UK and the MRC. The authors thank the participants, General Practitioners and staff of the EPIC-Norfolk study team, and collaborators for their contribution.

Generation Scotland: Chief Scientist Office of the Scottish Government Health Directorate (CZD/16/6), the Scottish Funding Council (HR03006), Medical Research Council UK, and a Wellcome Trust Strategic Award ‘STratifying Resilience and Depression Longitudinally’ (104036/Z/14/Z)(STRADL). Genotyping of the GS:SFHS samples was carried out by the Edinburgh Clinical Research Facility, University of Edinburgh. We are grateful to all the families who took part in the Generation Scotland: Scottish Family Health Study, the general practitioners and Scottish School of Primary Care for their help in recruiting them, and the whole Generation Scotland team, which includes academic researchers, IT staff, laboratory technicians, statisticians and research managers.

NFBC1966: received financial support from the Academy of Finland (project grants 104781, 120315, 129269, 1114194, 24300796, Center of Excellence in Complex Disease Genetics and SALVE), University Hospital Oulu, Biocenter, University of Oulu, Finland (75617), NHLBI grant 5R01HL087679- 02 through the STAMPEED program (1RL1MH083268-01), NIH/NIMH (5R01MH63706:02), ENGAGE project and grant agreement HEALTH-F4-2007-201413, EU FP7 EurHEALTHAgeing −277849, the Medical Research Council, UK (G0500539, G0600705, G1002319, PrevMetSyn/SALVE) and the MRC, Centenary Early Career Award. The program is currently being funded by the H2020-633595 DynaHEALTH action, academy of Finland EGEA-project (285547) and EU H2020 ALEC project (Grant Agreement 633212).

NFBC1986: received financial support from EU QLG1-CT-2000-01643 (EUROBLCS) Grant no. E51560, NorFA Grant no. 731, 20056, 30167, USA / NIHH 2000 G DF682 Grant no. 50945.

PIVUS: Swedish Research Council, Swedish Heart-Lung Foundation, Swedish Diabetes Foundation and Uppsala University. Genotyping and analysis was funded by the Wellcome Trust under awards WT064890 and WT098017. The PIVUS investigators express their deepest gratitude to the study participants.

SHIP / SHIP-Trend: Federal Ministry of Education and Research (01ZZ9603, 01ZZ0103, 01ZZ0403, 03ZIK012), and the German Research Foundation (GR 1912/5-1).

UKHLS: Economic and Social Research Council (ES/H029745/1). These data are from Understanding Society: The UK Household Longitudinal Study, which is led by the Institute for Social and Economic Research at the University of Essex and funded by the Economic and Social Research Council. The data were collected by NatCen and the genome wide scan data were analysed by the Wellcome Trust Sanger Institute. The Understanding Society DAC have an application system for genetics data and all use of the data should be approved by them. The application form is at: https://www.understandingsociety.ac.uk/about/health/data. A full list of authors is available in the Supplementary Note.

VIKING: Supported by a MRC Human Genetics Unit quinquennial programme grant “QTL in Health and Disease”. VIKING DNA extractions and genotyping were performed at the Edinburgh Clinical Research Facility, University of Edinburgh.

pQTL: The INTERVAL study: NHSBT (11-01-GEN) and the NIHR-BTRU in Donor Health and Genomics (NIHR BTRU-2014-10024) at the University of Cambridge in partnership with NHSBT. This study was partially funded by Merck. The views expressed are those of the authors and not necessarily those of the NHS, the NIHR, the Department of Health of England, or NHSBT. The Cardiovascular Epidemiology Unit at the University of Cambridge: UK MRC (G0800270), BHF (SP/09/002), UK NIHR Cambridge Biomedical Research Centre, ERC (268834), and European Commission Framework Programme 7 (HEALTH-F2-2012-279233).

### Competing interests

The following authors report potential competing interests:

L.V.W.: Louise V. Wain has received grant support from GSK.

M.D.T.: Martin D. Tobin has received grant support from GSK.

I.P.H.: Ian P. Hall has received support from GSK and BI.

K.S.: Kijoung Song is an employee of GlaxoSmithKline and may own company stock.

J.D.: John Danesh reports personal fees and non-financial support from Merck Sharp & Dohme (MSD) and Novartis, and grants from British Heart Foundation, European Research Council, MSD, NIHR, NHS Blood and Transplant, Novartis, Pfizer, UK MRC, Wellcome Trust, and AstraZeneca.

J.D.H.: Josh D. Hoffman is an employee of GlaxoSmithKline and may own company stock.

N.L.: Nick Locantore is an employee of GlaxoSmithKline and may own company stock.

J.C.M.: Joseph. C. Maranville was a Merck employee during this study, and is now a Celgene employee.

H.R.: Heiko Runz was a Merck employee during this study, and is now a Biogen employee.

R. T-S.: Ruth Tal-Singer is an employee of GlaxoSmithKline and owns company stock.

J.C.W.: John C. Whittaker is an employee of GlaxoSmithKline and may own company stock.

L.M.Y.: Laura M. Yerges Armstrong is an employee of GlaxoSmithKline and may own company stock.

E.K.S.: In the past three years, Edwin K. Silverman received honoraria from Novartis for Continuing Medical Education Seminars and grant and travel support from GlaxoSmithKline

A.S.B.: Adam S. Butterworth reports grants from Merck, Pfizer, Novartis, Biogen and Bioverativ and personal fees from Novartis

R.A.S.: Robert A. Scott is an employee of GlaxoSmithKline and may own company stock.

M.H.C.: Michael H. Cho has received grant support from GSK.

## References

1. Young, R.P., Hopkins, R. & Eaton, T.E. Forced expiratory volume in one second: not just a lung function test but a marker of premature death from all causes. Eur Respir J 30, 616–22 (2007).

2. Lefaudeux, D. et al. U-BIOPRED clinical adult asthma clusters linked to a subset of sputum omics. J Allergy Clin Immunol 139, 1797–1807 (2017).

3. Hekking, P.P. et al. Pathway discovery using transcriptomic profiles in adult-onset severe asthma. J Allergy Clin Immunol (2017).

4. Hobbs, B.D. et al. Genetic loci associated with chronic obstructive pulmonary disease overlap with loci for lung function and pulmonary fibrosis. Nat Genet 49, 426–432 (2017).

5. Salvi, S.S. & Barnes, P.J. Chronic obstructive pulmonary disease in non-smokers. Lancet 374, 733–43 (2009).

6. Nelson, M.R. et al. The support of human genetic evidence for approved drug indications. Nat Genet 47, 856–60 (2015).

7. Wilk, J.B. et al. A genome-wide association study of pulmonary function measures in the Framingham Heart Study. PLoS Genet 5, e1000429 (2009).

8. Repapi, E. et al. Genome-wide association study identifies five loci associated with lung function. Nat Genet 42, 36–44 (2010).

9. Hancock, D.B. et al. Meta-analyses of genome-wide association studies identify multiple loci associated with pulmonary function. Nat Genet 42, 45–52 (2010).

10. Soler Artigas, M. et al. Genome-wide association and large-scale follow up identifies 16 new loci influencing lung function. Nat Genet 43, 1082–90 (2011).

11. Cho, M.H. et al. A genome-wide association study of COPD identifies a susceptibility locus on chromosome 19q13. Hum Mol Genet 21, 947–57 (2012).

12. Loth, D.W. et al. Genome-wide association analysis identifies six new loci associated with forced vital capacity. 46, 669–77 (2014).

13. Wain, L.V. et al. Novel insights into the genetics of smoking behaviour, lung function, and chronic obstructive pulmonary disease (UK BiLEVE): a genetic association study in UK Biobank. Lancet Respir Med 3, 769–81 (2015).

14. Lutz, S.M. et al. A genome-wide association study identifies risk loci for spirometric measures among smokers of European and African ancestry. BMC Genet 16, 138 (2015).

15. Soler Artigas, M. et al. Sixteen new lung function signals identified through 1000 Genomes Project reference panel imputation. Nat Commun 6, 8658 (2015).

16. Hobbs, B.D. et al. Exome Array Analysis Identifies a Common Variant in IL27 Associated with Chronic Obstructive Pulmonary Disease. 194, 48–57 (2016).

17. Jackson, V. et al. Meta-analysis of exome array data identifies six novel genetic loci for lung function [version 1; referees: 1 approved with reservations]. Wellcome Open Research 3(2018).

18. Wain, L.V. et al. Genome-wide association analyses for lung function and chronic obstructive pulmonary disease identify new loci and potential druggable targets. Nat Genet 49, 416–425 (2017).

19. Wyss, A.B. et al. Multiethnic Meta-analysis Identifies New Loci for Pulmonary Function. bioRxiv (2017).

20. Loh, P.R. et al. Efficient Bayesian mixed-model analysis increases association power in large cohorts. Nat Genet 47, 284–90 (2015).

21. Benyamin, B. et al. GWAS of butyrylcholinesterase activity identifies four novel loci, independent effects within BCHE and secondary associations with metabolic risk factors. Hum Mol Genet 20, 4504–14 (2011).

22. Hammarsjo, A., Wang, Z., Vaz, R. & Taylan, F. Novel KIAA0753 mutations extend the phenotype of skeletal ciliopathies. 7, 15585 (2017).

23. Stephen, J. et al. Mutations in KIAA0753 cause Joubert syndrome associated with growth hormone deficiency. Hum Genet 136, 399–408 (2017).

24. Loukil, A., Tormanen, K. & Sütterlin, C. The daughter centriole controls ciliogenesis by regulating Neurl-4 localization at the centrosome. The Journal of Cell Biology (2017).

25. He, R. et al. LRRC45 is a centrosome linker component required for centrosome cohesion. Cell Rep 4, 1100–7 (2013).

26. Conkar, D. et al. The centriolar satellite protein CCDC66 interacts with CEP290 and functions in cilium formation and trafficking. 130, 1450–1462 (2017).

27. Uhlén, M. et al. Tissue-based map of the human proteome. Science 347(2015).

28. Hao, K. et al. Lung eQTLs to help reveal the molecular underpinnings of asthma. PLoS Genet 8, e1003029 (2012).

29. Lamontagne, M. et al. Refining susceptibility loci of chronic obstructive pulmonary disease with lung eqtls. PLoS One 8, e70220 (2013).

30. Obeidat, M. et al. GSTCD and INTS12 regulation and expression in the human lung. PLoS One 8, e74630 (2013).

31. Westra, H.J. et al. Systematic identification of trans eQTLs as putative drivers of known disease associations. Nat Genet 45, 1238–1243 (2013).

32. Consortium, G. Human genomics. The Genotype-Tissue Expression (GTEx) pilot analysis: multitissue gene regulation in humans. Science 348, 648–60 (2015).

33. Kundaje, A. et al. Integrative analysis of 111 reference human epigenomes. Nature 518, 317–30 (2015).

34. Sun, B.B. et al. Genomic atlas of the human plasma proteome. Nature 558, 73–79 (2018).

35. Dunham, I., Kulesha, E., Iotchkova, V., Morganella, S. & Birney, E. FORGE: A tool to discover cell specific enrichments of GWAS associated SNPs in regulatory regions. F1000Research 4(2015).

36. Zhou, J. & Troyanskaya, O.G. Predicting effects of noncoding variants with deep learning-based sequence model. Nat Methods 12, 931–4 (2015).

37. Cotto, K.C. et al. DGIdb 3.0: a redesign and expansion of the drug gene interaction database. Nucleic Acids Research, gkx1143–gkx1143 (2017).

38. Slack, R. et al. P112 Discovery of a Novel, High Affinity, Small Molecule αvβ6 Inhibitor for the Treatment of Idiopathic Pulmonary Fibrosis. QJM: An International Journal of Medicine 109, S60–S60 (2016).

39. Raab-Westphal, S., Marshall, J.F. & Goodman, S.L. Integrins as Therapeutic Targets: Successes and Cancers. Cancers (Basel) 9(2017).

40. Merrill, J.T. et al. Efficacy and Safety of Atacicept in Patients With Systemic Lupus Erythematosus: Results of a Twenty-Four-Week, Multicenter, Randomized, Double-Blind, Placebo-Controlled, Parallel-Arm, Phase IIb Study. Arthritis Rheumatol 70, 266–276 (2018).

41. Aschard, H. et al. Evidence for large-scale gene-by-smoking interaction effects on pulmonary function. Int J Epidemiol 46, 894–904 (2017).

42. Pulley, J.M. et al. Accelerating Precision Drug Development and Drug Repurposing by Leveraging Human Genetics. ASSAY and Drug Development Technologies 15, 113–119 (2017).

43. Yengo, L. et al. Meta-analysis of genome-wide association studies for height and body mass index in ∼700,000 individuals of European ancestry. bioRxiv (2018).

44. Wain, L.V. et al. Whole exome re-sequencing implicates CCDC38 and cilia structure and function in resistance to smoking related airflow obstruction. PLoS Genet 10, e1004314 (2014).

45. Black, P.N. et al. Changes in elastic fibres in the small airways and alveoli in COPD. Eur Respir J 31, 998–1004 (2008).

46. Løkke, A., Lange, P., Scharling, H., Fabricius, P. & Vestbo, J. Developing COPD: a 25 year follow up study of the general population. Thorax 61, 935–939 (2006).

47. Pulit, S.L., With, S.A.J. & Bakker, P.I.W. Resetting the bar: Statistical significance in whole-genome sequencing-based association studies of global populations. Genetic Epidemiology 41, 145–151 (2017).

48. Martinez, F.J. et al. A New Approach for Identifying Patients with Undiagnosed Chronic Obstructive Pulmonary Disease. Am J Respir Crit Care Med 195, 748–756 (2017).

49. Strug, L.J. et al. Cystic fibrosis gene modifier SLC26A9 modulates airway response to CFTR-directed therapeutics. Hum Mol Genet 25, 4590–4600 (2016).

50. Pulit, S.L., de With, S.A. & de Bakker, P.I. Resetting the bar: Statistical significance in whole-genome sequencing-based association studies of global populations. Genet Epidemiol 41, 145–151 (2017).

51. Bycroft, C. et al. Genome-wide genetic data on ∼500,000 UK Biobank participants. bioRxiv (2017).

52. Bulik-Sullivan, B.K. et al. LD Score regression distinguishes confounding from polygenicity in genome-wide association studies. Nature Genetics 47, 291 (2015).

53. The 1000 Genomes Project, C. A map of human genome variation from population-scale sequencing. Nature 467, 1061 (2010).

54. McCarthy, S. et al. A reference panel of 64,976 haplotypes for genotype imputation. Nature genetics 48, 1279–1283 (2016).

55. Yang, J., Lee, S.H., Goddard, M.E. & Visscher, P.M. GCTA: a tool for genome-wide complex trait analysis. Am J Hum Genet 88, 76–82 (2011).

56. Palmer, L.J. et al. Familial aggregation and heritability of adult lung function: results from the Busselton Health Study. Eur Respir J 17, 696–702 (2001).

57. Wilk, J.B. et al. Evidence for major genes influencing pulmonary function in the NHLBI family heart study. Genet Epidemiol 19, 81–94 (2000).

58. Wakefield, J. Reporting and interpretation in genome-wide association studies. Int J Epidemiol 37, 641–53 (2008).

59. van de Bunt, M., Cortes, A., Brown, M.A., Morris, A.P. & McCarthy, M.I. Evaluating the Performance of Fine-Mapping Strategies at Common Variant GWAS Loci. PLoS Genet 11, e1005535 (2015).

60. Wang, K., Li, M. & Hakonarson, H. ANNOVAR: functional annotation of genetic variants from high-throughput sequencing data. Nucleic Acids Res 38, e164 (2010).

61. Shihab, H.A. et al. Predicting the functional, molecular, and phenotypic consequences of amino acid substitutions using hidden Markov models. Hum Mutat 34, 57–65 (2013).

62. Jansen, R. et al. Conditional eQTL analysis reveals allelic heterogeneity of gene expression. Hum Mol Genet 26, 1444–1451 (2017).

63. Battle, A., Brown, C.D., Engelhardt, B.E. & Montgomery, S.B. Genetic effects on gene expression across human tissues. Nature 550, 204–213 (2017).

64. Consortium, E.P. et al. An integrated encyclopedia of DNA elements in the human genome. Nature 489, 57–74 (2012).

65. MacArthur, J. et al. The new NHGRI-EBI Catalog of published genome-wide association studies (GWAS Catalog). Nucleic Acids Research 45, D896–D901 (2017).

66. Leslie, R., O’Donnell, C.J. & Johnson, A.D. GRASP: analysis of genotype-phenotype results from 1390 genome-wide association studies and corresponding open access database. Bioinformatics 30, i185–94 (2014).

